# Reasoning with programs in replay

**DOI:** 10.1101/2025.10.10.681671

**Authors:** Sebastijan Veselic, Nour Mohsen, Lennart Luettgau, Elena Gutierrez, Maria K. Eckstein, Steven W. Kennerley, Timothy E.J. Behrens, Timothy H. Muller, Zeb Kurth-Nelson

## Abstract

Reasoning flexibly composes known elements to solve novel problems. Recent theories suggest the brain uses the axis of time to compose elements for reasoning. In this view, elements are packaged into fast neural sequences, with each sequence exploring the implications of a different composition. Using magnetoencephalography, we tested this idea while participants mentally executed programs. Each program contained a set of steps linked to operations, the implications of which had to be computed. We found that behavioral performance scaled with program complexity, and by the end of execution, inferred program solutions were represented in prefrontal and parietal cortices. We identified a possible mechanism by which these solutions were computed. During reasoning, representations of steps in the program reactivated in fast sequences, consistent with sampling candidate partial solutions. Further evidence suggested these reactivations were accompanied by representations of their operations, and were followed by neural patterns reflecting their computed implications. Together, these results suggest replaying sequences supports program execution and reveal a highly organized temporal microarchitecture of reasoning.

## 1 Introduction

Humans have the remarkable ability to flexibly execute mental operations when reasoning about new problems. If we know how to multiply, we can use this operation to solve equations during exams or to calculate dinner ingredients for ten people instead of one. Reasoning requires composing existing representations in novel ways. Recent empirical and theoretical work suggests replay as a candidate neural mechanism supporting this flexible composition, and thereby a mechanism for general-purpose reasoning^1–4^.

Replay was first discovered in hippocampal place cells. During sleep, these cells spontaneously reactivated the same sequences of locations the animal had visited while awake^5–8^. Early theories proposed that replay rehearsed experience to etch into long-term memory^9–12^. However, over the last 30 years, evidence has accumulated for a more general function of replay.

A robust observation is that replay can synthesize new trajectories rather than simply recalling past ones. For example, if two adjacent sections of a maze are experienced separately, replay spontaneously generates new sequences in which both sections are joined together to span the entire path^13^. Similarly, when rodents visually observe a novel environment, replay starts playing out paths in that environment before the animal has taken them^14;15^. It is now widely accepted that replay sequences are drawn from an internal world model or cognitive map^16–18^.

From human replay experiments using magnetoencephalography (MEG), we have additionally learned that replay is not limited to locations in physical space. When participants learn about a set of objects arranged in a graph with non-spatial topology, the brain spontaneously replays paths through that graph^19^. If participants then learn an abstract rule that changes the graph’s topology, replay sequences rapidly reorganize to reflect the new topology^1^.

Moreover, replay can include abstract representations of an object’s position in a sequence^1^. These representations are abstract because they generalize across different sequences. For example, any object appearing in position 2 shares the abstract ‘position 2’ representation, regardless of which sequence it belongs to. As sequences of objects are replayed, each object representation is accompanied by a reactivation of its position in tight temporal coordination. More recently, Schwartenbeck et al.^2^ found suggestive evidence that replay sequences are not constrained to follow paths along a Markov Decision Process (MDP). In their experiment, participants were trained to solve combinatorial puzzles. Replay sequences were composed of puzzle pieces such that each sequence represented a possible configuration of the puzzle. Thus, rather than only forming a series of consecutive locations, replay might encode structured information like ‘Lucy eats watermelon’ by playing out ‘Lucy’, ‘eats’, and ‘watermelon’ as a sequence.

In response to the evolving empirical data, we proposed a theory of replay as a general-purpose compositional reasoning mechanism^3^. In this theory, a replay sequence is a set of entities composed together into a larger compositional structure, where each entity is accompanied by an abstract representation of its role within the structure. Roles describe how entities fit together, enabling reasoning about the meaning of the structure as a whole. For example, in the sentence ‘Lucy eats watermelon’, the grammatical role of ‘Lucy’ is ‘subject’. Knowing that Lucy is the *subject* enables computing the implications of ‘Lucy eats watermelon’, such as the subsequent contents of Lucy’s stomach.

To test predictions of this theory, we designed a reasoning task in which participants mentally executed a program on each trial (Figure 1). We provide behavioral and neural evidence for program execution, with a representation of the inferred program solution emerging in prefrontal and parietal cortices by the end of execution. During execution, we demonstrate locations in programs were replayed in fast neural sequences, that these were temporally aligned with representations of their associated roles, as well as their implications. This reveals a highly-organized temporal micro-architecture during reasoning.

**Figure 1:**
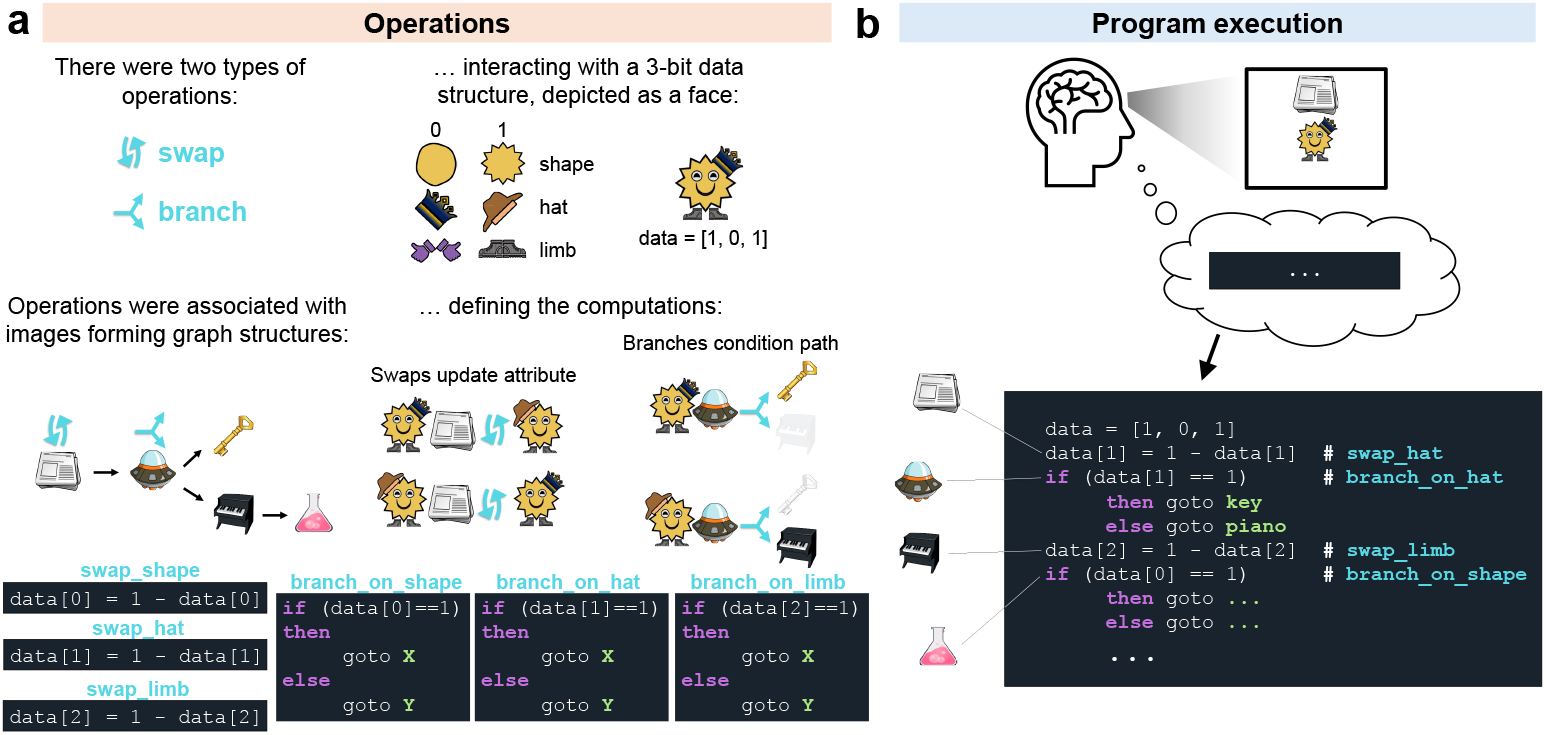
Conceptual overview of the study: operations and program execution. a) Participants learned about *swap* and *branch* operations, acting as roles in our task. Each operation was associated with an image object such as Newspaper, Spaceship, Piano, or Key, representing locations in a graph. Paths through this graph were “programs” that interacted with a cartoon face made up of three binary attributes: shape (round or spiky), hat (crown or fedora), and limb (hands or feet). Note that symbols for branch and swap are only included for illustration and were not shown to participants. b) On each trial, participants mentally executed a novel program: they were given the location (e.g., Newspaper) and face (e.g., [Spiky, Crown, Feet]) from which to start executing a sequence of operations.

## 2 Results

Before executing programs in the reasoning task during MEG recording, participants first learned the necessary task components over two days of training (see Methods: Training: Day 1 & Day 2 and Testing: Day 3).

On Day 1, participants learned a directed graph of 12 program locations (displayed as image objects) plus one END token (Figure 2A). On Day 2, participants learned that each program location was associated with an operation. There were six operations: swap shape, swap limb type, swap hat, branch on shape, branch on limb type and branch on hat. Each operation appeared twice in the graph and interacted with a cartoon face data structure (Figure 2B). Participants received no advance information about the Day 3 task during Day 1 & Day 2 training.

**Figure 2:**
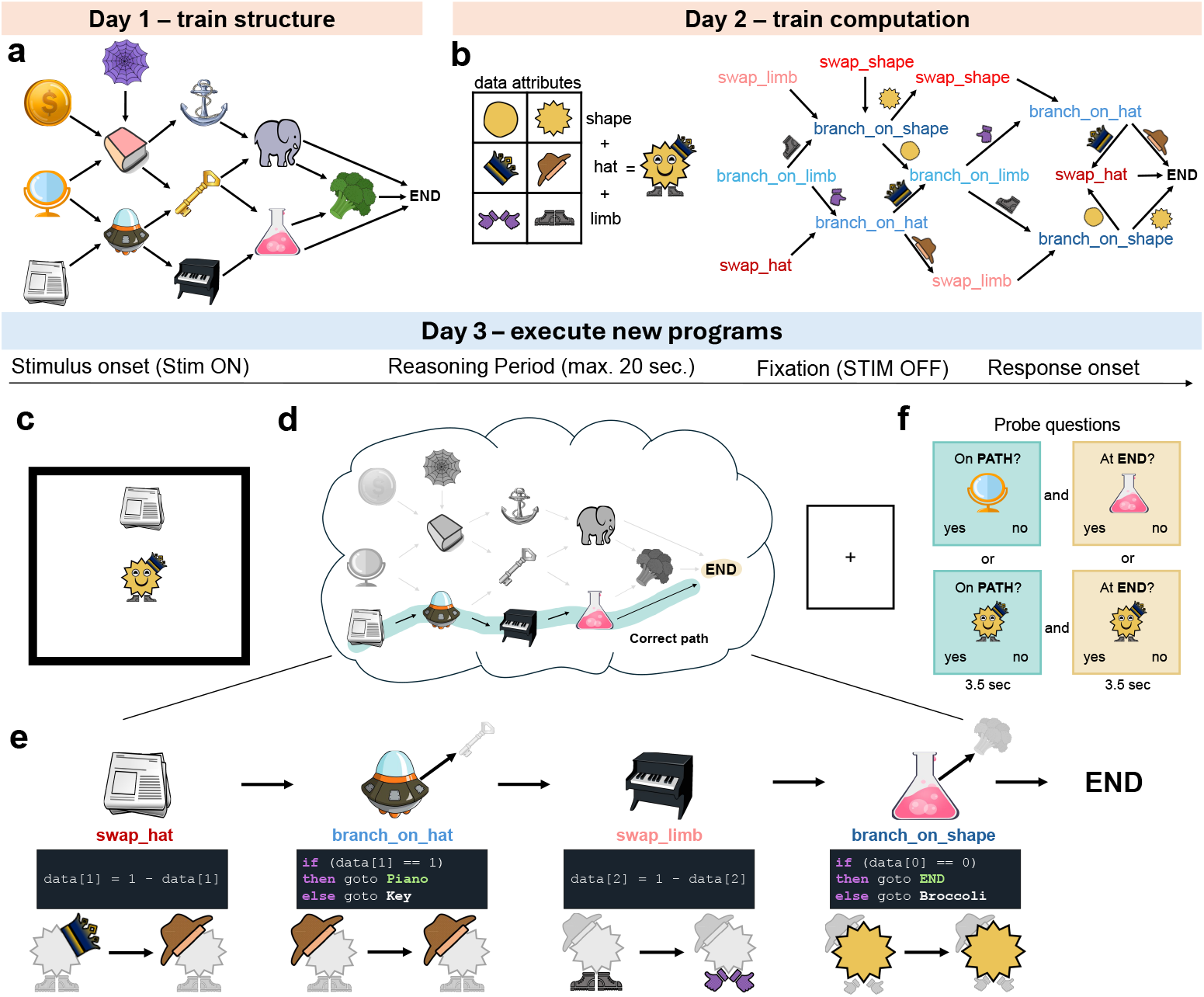
Study design and task description. **a)** On Day 1, participants learned all possible transitions between image objects. Each participant was randomly assigned one of four graphs; only one is shown. Note that participants never explicitly saw the graph. **b)** On Day 2, participants learned cartoon face attributes and program operations (“operations”). Left panel: cartoon faces were generated from all combinations of shape, limb, and hat. Right panel: Each object from Day 1 was associated with an operation. For example, the Newspaper location required a swap hat operation while Spaceship required a branch on hat operation. By the end of Day 2, participants learned that each object was a “location” in a program graph. Branches transitioned to two possible program locations. Swaps transitioned to one. **c)** Day 3 reasoning task stimulus onset. A starting program location and face remained on-screen during the reasoning period. **d)** During reasoning, participants had to mentally execute the program implied by the pairing until reaching the END token within 20 seconds. **e)** Program execution schematic for the trial from panel c. Note that irrelevant attributes at each operation are grayed out. **f)** After reasoning, two probe questions (END, PATH) were presented serially for 3.5 seconds per question in a randomized order. Within each trial, both probe questions referred to either objects or faces, randomized across trials.

Day 3 (“Reasoning Task”) took place in the MEG scanner. On each trial, participants were shown a random pairing of one program location (object) and one cartoon face (Figure 2C). Both stimuli remained on-screen for 20 seconds. During this time participants had to mentally execute the program defined by that pairing until reaching the END token (“Reasoning Period”, Figure 2D). Program execution consisted of performing the operation associated with each program location (Figure 2E). At swap operations, this meant mentally updating the relevant face attribute and mentally moving to the next program location. For example, starting with Newspaper and Crown required updating the hat from Crown to Fedora (Crown → Fedora) and moving to Spaceship. At branch operations, the value of the relevant face attribute determined the next program location. For example, when Spaceship was paired with Fedora, the next program location was Piano, otherwise the next location would have been Key. In this case, no data update was required. Programs were deterministic. Therefore, each trial had one correct execution path (“Correct path”) which consisted of three or four operations. Participants were allowed to terminate the reasoning period early by pressing a button. Crucially, the programs on Day 3 were novel and no feedback was provided during reasoning; programs were executed mentally.

After the reasoning period, participants answered two probe questions on every trial. Probes were designed to test whether they had correctly executed the program (Figure 2F). PATH questions (Figure 2F, green highlight) asked about an object or face that was encountered before the final step of execution. END questions (Figure 2F, orange highlight) asked about the final object or face in their execution path (see also Methods: Reasoning Task).

### 2.1 Behavioral evidence for program execution

We were first interested in whether the behavioral data were consistent with participants mentally executing programs. Participants’ behavioral accuracy on probe questions was above chance on all questions (all p <.001) and higher on questions about objects compared to faces (END: *t*_28_ = 2.74, p =.011, PATH: *t*_28_ = 5.78, p <.001, Figure 3A). Across participants, END and PATH accuracy were correlated (Figure 3B). Within participants, a correct answer to the END question on a given trial predicted a correct answer to the PATH question on the same trial (*t*_28_ = 2.17, p =.039, Figure 3C), consistent with participants reaching the END token by executing the program up to that point.

**Figure 3:**
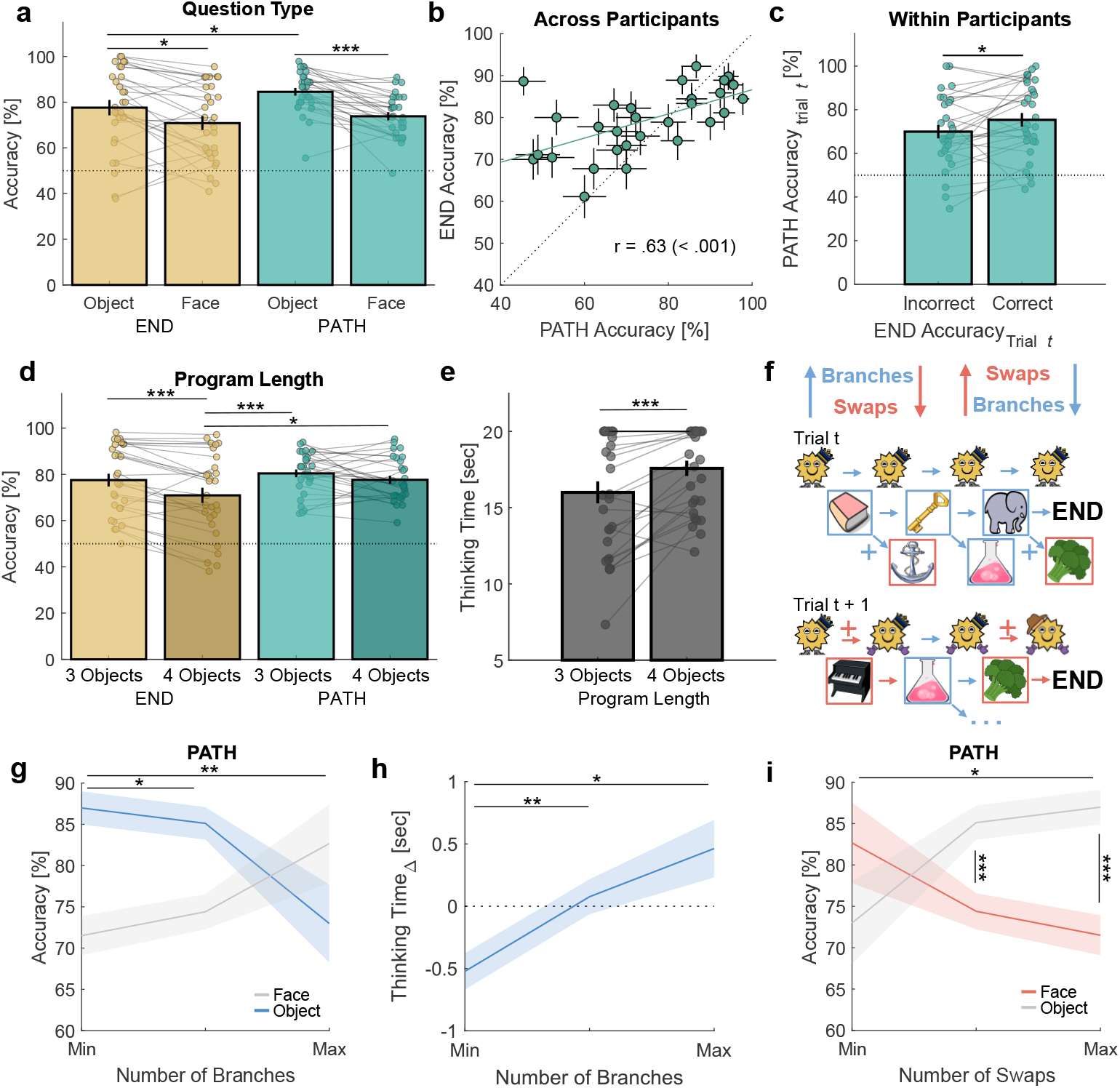
Behavior is consistent with mental execution of programs. a) Accuracy across question type (END, PATH) split by modality (Object, Face). Dotted line represents chance accuracy of accepting / rejecting the probe question on a trial-by-trial basis (see Methods: Behavioral analyses). Each dot represents a participant. b) END and PATH accuracy correlated across participants. Error bars represent standard error of the mean (SEM) within participant. Dotted diagonal line represents equal performance across both question types. c) On trials where END questions were answered correctly, PATH questions were also more likely to be answered correctly. d) END accuracy was lower on longer programs. e) Thinking time was longer on longer programs. f) More branches in the correct program required more objects to be evaluated. This increased memory load for PATH Object questions but decreased it for PATH Face questions (because fewer or no face swaps were required). The opposite symmetry is true for trials with more swaps in the correct program. g) PATH Object trial accuracy (blue shading) decreased as number of branches increased. The gray shading depicts accuracy on PATH Face trials. h) Participants’ thinking time increased as the number of branches in the correct program increased. i) PATH Face trial accuracy (red shading) decreased as the number of swaps increased. The gray shading depicts accuracy on PATH Object trials. Thick lines represent mean accuracy across participants, error bars represent SEM across participants. *** <.001, ** <.01, * <.05.

If participants were mentally executing programs, then accuracy and thinking time should be sensitive to program length and content. As program length increased, accuracy decreased (*t*_28_ = 4.53, p <.001, Figure 3D). Furthermore, END accuracy was lower than PATH accuracy on the longest programs, even though no feedback was provided during reasoning (*t*_28_ = 2.33, p = 0.027. See also Supplementary Figure 2A). In addition, participants’ thinking time increased on longer programs (*t*_28_ = 4.13, p <.001, Figure 3E), analogous to Jensen et al.^20^ where participants’ thinking time increased as planning trajectory length increased.

In addition to program length, program content also varied across trials (Figure 3F). When the number of branches in a program increased, this increased the potential number of program locations to be evaluated. Correspondingly, we found PATH Object accuracy decreased as the number of branches increased (*t*_28_ = 3.16, p =.004, Figure 3G blue shading. See also Supplementary Figure 2B). Furthermore, as the number of branches increased, thinking time was longer even after controlling for program length (*t*_28_ = 2.89, p =.008, Figure 3H. See also Supplementary Figure 2C). In contrast, as the number of swaps increased, Face accuracy decreased (PATH: *t*_28_ = 2.09, p =.046, Figure 3I red shading. END: *t*_28_ = 2.96, p =.006, Supplementary Figure 2B). Because an operation was either a branch or a swap, programs with more branches had fewer swaps and vice versa. Thus, as the number of branches increased, PATH Face accuracy increased (Figure 3G gray shading), and as the number of swaps increased, PATH Object increased (Figure 3I gray shading). The same data are visualized twice across both panels to highlight the symmetry between branches and swaps.

Overall, the behavioral data were consistent with participants mentally executing programs during the reasoning period.

### 2.2 Neural representations of program paths emerge during reasoning

We next examined whether participants’ neural activity was related to the programs they had to execute. To answer probe questions, participants in principle needed to retain each trial’s execution path in working memory. Thus, if participants executed programs, trials with more similar paths should also produce more similar neural patterns by the end of the reasoning period. To test this, we computed the trial-by-trial program similarity, defined as the number of overlapping locations in the correct execution paths between trials (Figure 4A. See Methods: RSA analyses). We then used representational similarity analysis (RSA) and tested whether program similarity predicted neural activity. Because thinking time varied across trials and participants, we focused on the start and end of the reasoning period.

**Figure 4:**
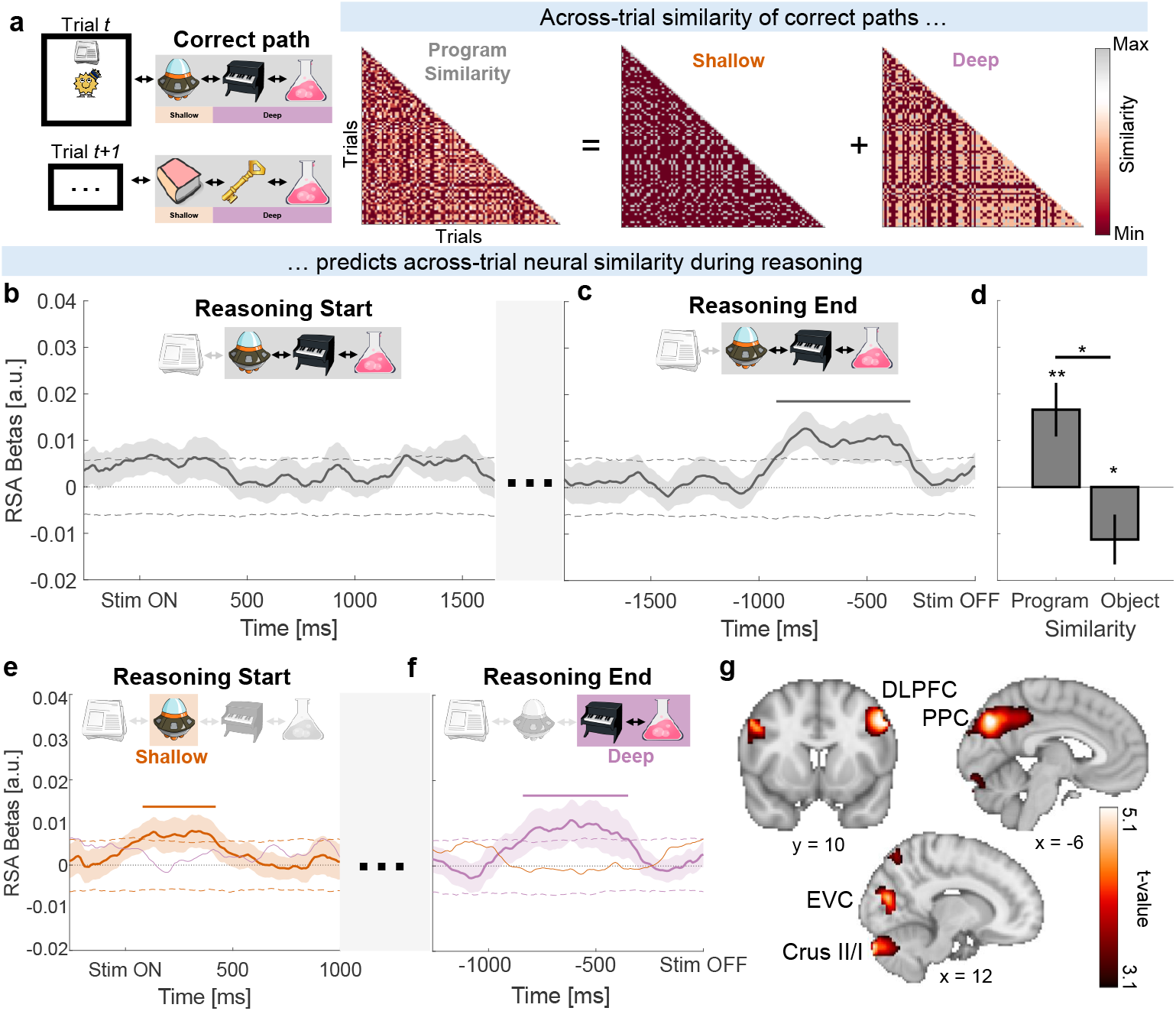
Across-trial program similarity predicts neural activity during reasoning. a) Program similarity is the position-dependent overlap of program locations on the *inferred* correct path across trials (see Methods: RSA analyses). Program similarity comprised Shallow and Deep locations. Shallow locations were the associations of the starting location, i.e. first location *not* shown on the screen. Deep locations were associations of Shallow locations. b) At Reasoning Start, program similarity (RSA betas) did not predict neural activity. Dashed lines represent the 97.5th percentile of a null distribution for visual purposes. c) At Reasoning End, program similarity predicted participants’ neural activity. The black bar represents significance in cluster-based permutation testing (exceeding the 97.5th percentile of the mass-corrected null). d) During the last second of reasoning, program similarity (“Program”) rather than overlap of objects (“Object”) predicted participants’ neural activity (see Methods: RSA analyses for definitions). e) At Reasoning Start, Shallow program similarity (orange) predicted participants’ neural activity. The purple line represents Deep program similarity. f) At Reasoning End, the Deep program similarity (Purple) drove the effect in panel c. The orange line represents Shallow program similarity. g) Source reconstruction of the effect in panel c with clusters in DLPFC, PPC, EVC, and cerebellum (Crus II/I) that survived whole-brain cluster correction, FWE p <.05. ** p <.01, * <.05. Mean values and thick lines represent the average across participants, shading and error bars represent the SEM across participants.

As expected, program similarity did not predict neural activity at the start of the reasoning period (Figure 4B): given the task difficulty and program novelty, it would not have been plausible for participants to have consistently reasoned through the correct path at that point. By the end of the reasoning period, however, program similarity predicted neural activity (Figure 4C). This effect was specifically driven by program similarity - which measured the similarity of *paths* - rather than being driven only by the overlap of objects on those paths (Program: *t*_28_ = 2.88, p =.008, Program vs. Object: *t*_28_ = 2.67, p =.016, Figure 4D) when examining the last second of the reasoning period (see Methods: RSA analyses).

We next asked which program locations contributed to this effect. Program similarity was separated into “shallow” (Figure 4A orange shading) and “deep” (Figure 4A purple shading) locations. Within the first second of reasoning, shallow program similarity, in fact, predicted participants’ neural activity (Figure 4E). Given the short latency, this likely reflected learned associations between the starting locations and their immediate transitions^21^. In contrast, deep program similarity only emerged toward the end of reasoning (Figure 4F), consistent with participants progressing into deeper parts of the program.

Finally, we localized the program similarity signal from Figure 4C using beamforming of wideband neural activity (see Methods: RSA source Reconstruction)^1;22^. This revealed a network (Figure 4G) that included dorsolateral prefrontal cortex (DLPFC), posterior parietal cortex (PPC), cerebellum (Crus II/I), early visual cortex (EVC), and lateral occipital cortex. These regions have been linked to the maintenance and manipulation of information in working memory^23;24^, suggesting that program execution may have recruited these processes.

Overall, these results indicate that during the reasoning period, neural activity became increasingly related to program paths, with this information being maintained in working memory.

### 2.3 Replay of program locations

Considering that both the behavioral and neural evidence suggested participants executed programs, we next asked whether locations within those programs were replayed in fast neural sequences during the reasoning period^2;19;22^. To test this, we trained separate decoding models to identify neural representations of each object - indexing these locations using functional localizer data (Figure 5A-C. See Methods: functional localizer task and decoding). We then applied these decoders to unlabeled data from the reasoning period to detect spontaneous reactivations of objects, and used temporally delayed linear modeling (TDLM) to assess the extent to which sequences of program locations adhered to paths on the graph^1;19;22;25^.

**Figure 5:**
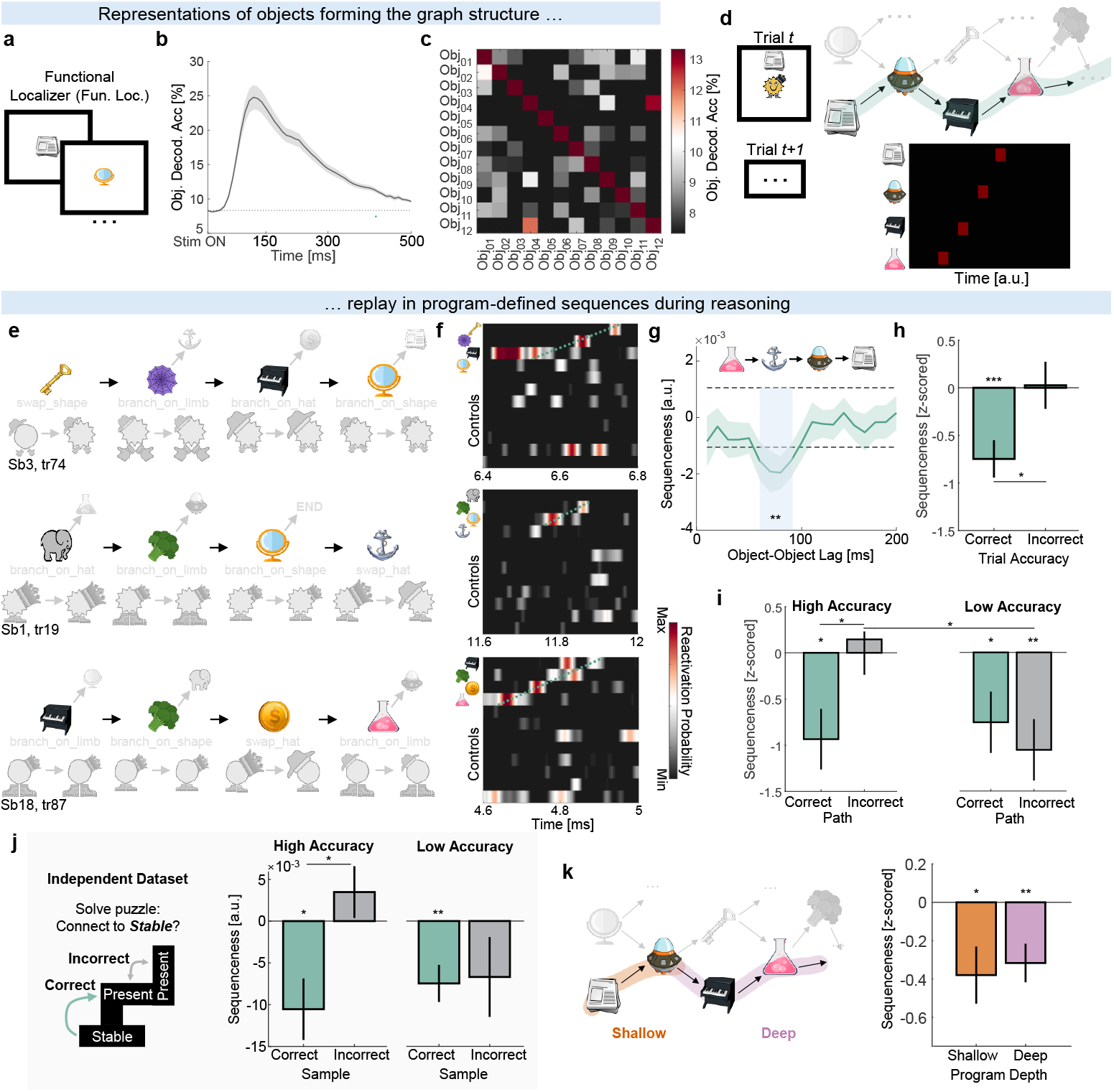
Correct and incorrect program paths were replayed in sequences during reasoning. a) Before the reasoning task, participants performed a functional localizer task in which objects were presented in randomized order (see Methods: Functional localizer task). b) Average cross-validated decoding accuracy for objects in the functional localizer. The dotted line represents chance (8.3%). c) Object decoding confusion matrix at the decoding peak (120 ms after stimulus onset). d) Each trial contained one correct path through the graph (green shading) that was defined by the starting location and face (see Figure 2D). All other paths were considered incorrect. We predicted program locations along the correct path would be replayed in sequences ^2;19;22;27^. e) Example program execution schematics for different trials and participants. Only program location reactivations were evaluated in this figure. f) Example correct paths and control object reactivations during the reasoning period from the program execution schematics in panel e. g) Correct path sequenceness during the reasoning period. Dashed lines represent the 95th percentile of the null distribution (multiple-comparisons corrected). h) Replay effect from panel g, split by probe question accuracy after reasoning. i) Replay effect from panel g for correct and incorrect paths, further split by participants’ overall probe accuracy (median split). See caption in panel d, Supplementary Figure 3A, and Methods: Location → Object sequences for definitions. j) Left panel: Conceptual summary of task from Schwartenbeck et al. ^2^. Participants were sampling between puzzle pieces that would connect to a *Stable* puzzle piece. Samples between a *Present* and *Stable* piece were considered correct, while samples between two *Present* pieces were considered incorrect. Right panel: Replication of panel i using data from Schwartenbeck et al. ^2^. Note that sequenceness values were inverted to match interpretation across datasets because the direction of sequenceness in their data were arbitrary (see Methods: Independent dataset analysis). k) Left panel: Distinction between Shallow (orange) and Deep (purple) program locations (see also Methods: RSA Analyses). Right panel: Replay effect from panel g, split by the depth of the program location on the correct path. *** p <.001, ** p <.01, * <.05. Mean values and thick lines represent the average across participants, shading and error bars represent the SEM across participants.

We first examined correct paths (Figure 5D green shading. See Figure 5E, F for examples) and found program locations were replayed with an ∼80 ms object-to-object lag (*t*_28_ = 3.14, *p* =.004, Figure 5G). This effect exceeded a permutation threshold corrected for multiple comparisons (see Methods: Sequenceness analyses) and was present only on trials where participants correctly answered probe questions *after* the reasoning period (Correct: *t*_28_ = 3.81, *p* <.001, Correct vs. Incorrect: *t*_26_ = 2.46, *p* =.021, Figure 5H). Importantly, the effect was present across participants when split by overall accuracy (High Accuracy: *t*_13_ = 2.85, *p* =.014, Low Accuracy: *t*_14_ = 2.28, *p* =.039, Figure 5I green shading).

We also examined incorrect paths (see Supplementary Figure 3A for diagram). We reasoned that participants may have sometimes executed incorrect programs due to uncertainty or error. In line with this, high-accuracy participants showed no replay of incorrect paths (Incorrect: *p* =.71, Correct vs. Incorrect: *t*_13_ = 2.70, *p* =.018). In contrast, low-accuracy participants additionally replayed incorrect paths (*t*_14_ = 3.18, *p* =.007, Figure 5I gray shading) and did so more compared to high-accuracy participants (*t*_27_ = 2.38, *p* =.024, Figure 5I gray shading).

Sampling of both correct and incorrect solutions during a thinking period was also observed by Schwartenbeck et al.^2^. Given our findings in the previous paragraph, we wondered if the same pattern might exist in the Schwartenbeck et al. data (see Methods: Independent dataset analysis). Indeed, re-analyzing their data revealed that although all participants replayed correct paths (High: *t*_9_ = 2.95, *p* =.016, Low: *t*_9_ = 3.55, *p* =.006), high-accuracy participants did not replay incorrect paths (Correct vs. Incorrect: *t*_9_ = 2.65, *p* =.026) with a significant interaction between accuracy and path type (*F*_1,18_ = 6.98, *p* = 0.017, Figure 5J). Our own data additionally align with rodent studies in which lower-performing animals show more reverse replay^26^. Consistent with this, participants with more reverse replay in our data also spent more time thinking during the reasoning period (r = -.49, p =.008. Supplementary Figure 3B), even after controlling for overall accuracy and correct path replay.

Finally, we asked whether replay of correct paths was driven by the first transition in the program, considering that the starting location remained on-screen during the reasoning period. We therefore repeated the TDLM analysis using separate regressors for shallow and deep program locations and found independent evidence for replay of both parts (Shallow: *t*_28_ = 2.18, *p* =.038, Deep: *t*_28_ = 2.91, *p* =.007, Figure 5K). In fact, we found a pattern suggesting participants replayed increasingly deeper program locations as the session progressed, paralleling improvements in behavioral performance (Supplementary Figure 3D, E).

### 2.4 Program locations are accompanied by their operations

Having found replay of program locations, we next asked whether a reactivation of a program location was accompanied by a reactivation of the operation at that location. We tested for such a pattern across four analyses. To do this, we trained decoders to detect representations of operations using the operation localizer data (Figure 6A. See Methods: operation localizer task). In this task, participants were shown an object and asked about the operation associated with that program location. Performance on this task was high (93.88% *±* 1.01 SEM) and correlated with reasoning task performance (r =.52, p =.005, Figure 6B. See also Supplementary Figure 4A).

**Figure 6:**
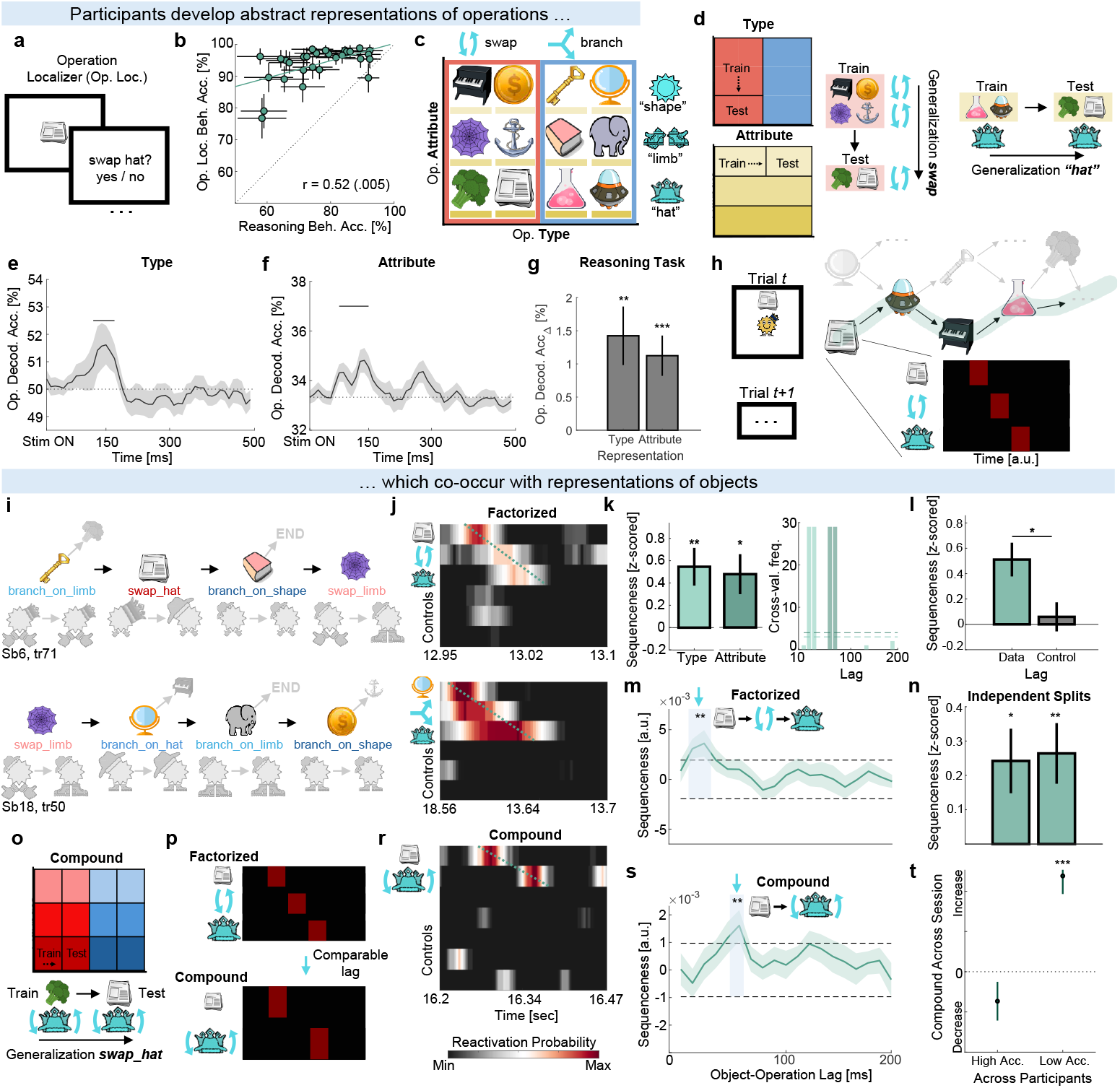
Reactivations of program locations are accompanied by reactivations of factorized and compound representations of operations. a) Operation Localizer. After the reasoning task, participants were shown objects and were told to recall their operation. They were required to accept / reject the subsequent text label on every trial (see Methods: Operation localizer task). b) Correlation between operation localizer and reasoning task accuracy. Each dot represents a participant. Error bars represent SEM within participant across both tasks. c) Operations factorized along two dimensions. On the x-axis: Type (six objects swap - red highlight, six objects branch - blue highlight). On the y-axis: Attribute (four objects use “shape”, four use “limb”, and four use “hat” - yellow highlights). Quotation marks for “shape” etc. emphasize Attribute representations are abstract and do not refer to specific face attribute values. d) Cross-validation scheme: Type and Attribute decoders had to generalize on held-out objects sharing an operation dimension without any overlap across dimensions. Such a train-test split ensured representations for Type and Attribute were orthogonal. e) Average cross-validated decoding accuracy for Type. Shading represents SEM across participants. The black bar represents significance in cluster-based permutation testing (exceeding the 97.5th percentile of the mass-corrected null). The dotted line represents chance (50%). f) Same as in panel e but for Attribute. g) Same as in panels e, f but taken from the reasoning task during object representations. Time points used for averaging were determined using a leave-one-participant-out cross-validation procedure (see Methods: Abstract representation decoding). h) On each trial of the reasoning task, we tested whether reactivations of program locations on the correct path were accompanied by reactivations of their operations. Note that all locations on the correct path were investigated. i) Example program execution schematics for different trials and participants with highlighted operations. j) Example Location Type → Attribute → and control reactivations. k) Left panel: Location → Type and Location → Attribute sequences estimated using a leave-one-participant-out cross-validation procedure (see Methods: Location → Operation sequences). Right panel: distribution of lags used for cross-validating the effects in the left panel. Dashed lines represent the 95th percentile of the null distributions (multiple-comparisons corrected). l) Type and Attribute sequences were stronger at lags observed in the data (30ms and 60ms) compared to control lags (60ms and 30 ms). m) Sequenceness for Location → Type → Attribute sequences. The effect in panel m is significant when estimated on independent splits (see Methods: Location Operation sequences). The cross-validation scheme for operations used the same logic as described in panel d and used in Liu et al. ^1^. p) We expected the factorized and compound sequences to have comparable lags. r) Example Location → Operation and control reactivations. s) Sequenceness for Location → Operation sequences. t) Beta coefficients of a trial-index regressor capturing the evolution of the effect in panel s across the session, shown separately for the High and Low accuracy participant group (median split) (see Methods: Location → Operation sequences). Error bars represent the SEM of this regressor. *** p <.001, ** p <.01, * <.05.

To isolate neural responses that would reflect representations of operations rather than object identity, we factorized operations into the two dimensions that define them (Figure 6C): Type (swap, branch) and Attribute (hat, limb, shape); in close analogy to Liu et al.^1^. We then followed the approach from Liu et al.^1^ and trained decoders using a cross-validation scheme where we held out all trials on which a certain object was shown (Figure 6D). For example, a decoder trained on swap shape (e.g. Piano, Coin) and swap limb (e.g. Web, Anchor) objects had to classify swap hat objects (e.g. Broccoli, Newspaper) as belonging to the swap category. Thus, decoders could only perform above chance if neural activity generalized across objects and reflected abstract Type or Attribute representations. Because object assignments were randomized across participants, performance could not be explained by shared visual features.

Even under this challenging cross-validation procedure, both Type and Attribute were decoded above chance on the localizer data (Figure 6E, F. See also Supplementary Figure 4B). As further validation, we tested these decoders on object presentations from reasoning task data (see Methods: Abstract representation decoding) which similarly revealed above-chance decoding (Type: *t*_28_ = 3.23, *p* =.003, Attribute: *t*_28_ = 3.70, *p* <.001, Figure 6G). We then applied object and operation decoders to the reasoning period and used TDLM to measure the extent to which their reactivations occurred in tight temporal coordination (Figure 6H. See Methods: Location → Operation sequences. See also Supplementary Figure 4C, D for validation simulations).

We first looked for reactivations of each operation dimension separately across two independent analyses. We found reactivations of program locations were accompanied by reactivations of Type and Attribute with a 10-30 ms and a 60 ms lag respectively (Figure 6I-L. See also Supplementary Figure 4E). These results suggest the presence of sequences containing factorized representations where locations (e.g., Newspaper) are accompanied by both their Type (swap) and Attribute (hat). This closely matches previous reports of factorized sequences occurring at comparable lags (40-60 ms)^1^.

Because the first two analyses isolated each dimension separately, we were able to combine both in a single design matrix based on the identified lags [Location → Type → Attribute]. In the third analysis, this approach confirmed the predicted effect where individual representations occurred with a 20–40 ms lag relative to one another (Figure 6M. See also Supplementary Figure 4F). Within participants, the effect was significant in correct trials (p <. 05) but not in incorrect trials (p >.21). Because this analysis was not independent of the previous two, we did an additional test of robustness. We split trials into odd and even halves. We repeated the first two analyses on odd trials, recorded their significant lags, and used these lags to repeat the third analysis on even trials. The effect survived in both independent cross-splits (Figure 6N).

Jointly, the first three analyses suggest sequences of program locations were accompanied by representations of their type and attribute, which together define a specific operation (e.g., swap hat). In the last analysis, we tested for the presence of “compound” sequences where program locations would be accompanied by representations of these specific operations themselves, rather than their factorized dimensions. We were able to conduct this analysis because each operation (e.g. swap hat) was associated with two objects (e.g., Broccoli and Newspaper). We repeated the cross-validation procedure described above by training on one object and testing on the other to ensure operation representations were not driven by object identity (Figure 6O). This approach additionally ensured this analysis was independent of the previous ones (see Methods: Location → Operation sequences). Because *swap* and *hat* together convey identical information as *swap hat*, we expected compound sequences would have lags similar to the factorized ones (Figure 6P). Indeed, as expected from the previous analyses, reactivations of program locations were also accompanied by specific operation reactivations at a ∼ 60 ms lag, matching the factorized pattern (Figure 6R, S). Similar to the third analysis, the effect was absent on incorrect trials (*p* =.97; Correct vs. Incorrect: *t*_26_ = 2.05, *p* =.049), suggesting that reactivations of program locations and their operations may have been involved in program execution.

These four analyses provide evidence that spontaneous reactivations of program locations were accompanied by operation Type (∼3 0 ms) and Attribute (∼ 60 ms), with the specific operation also reactivating at a comparable ∼ 60 ms lag: the first moment when both dimensions uniquely identify it. These results support the hypothesis that spontaneous reactivations of entities, likely as part of replay sequences, are bound to arbitrary roles^1;3^.

### 2.5 Results of reasoning appear immediately after replay events

If the information represented in spontaneous reactivations of program locations and operations is used for reasoning about implications, it is reasonable to ask whether the results of reasoning (updates of face attributes) appear in the brain immediately afterward. This was the question behind our final set of analyses.

Because operations interacted with the face data structure, we trained decoders to decode the two possible values of each attribute using functional localizer data (Figure 7A). Crossvalidation accuracy on the functional localizer was high (shape: 56%, limb: 64%, hat: 62%, Figure 7B), the models successfully decoded starting face attributes on the reasoning task data (Figure 7C), and decoded them in a manner that was consistent with starting operations (Supplementary Figure 5A).

**Figure 7:**
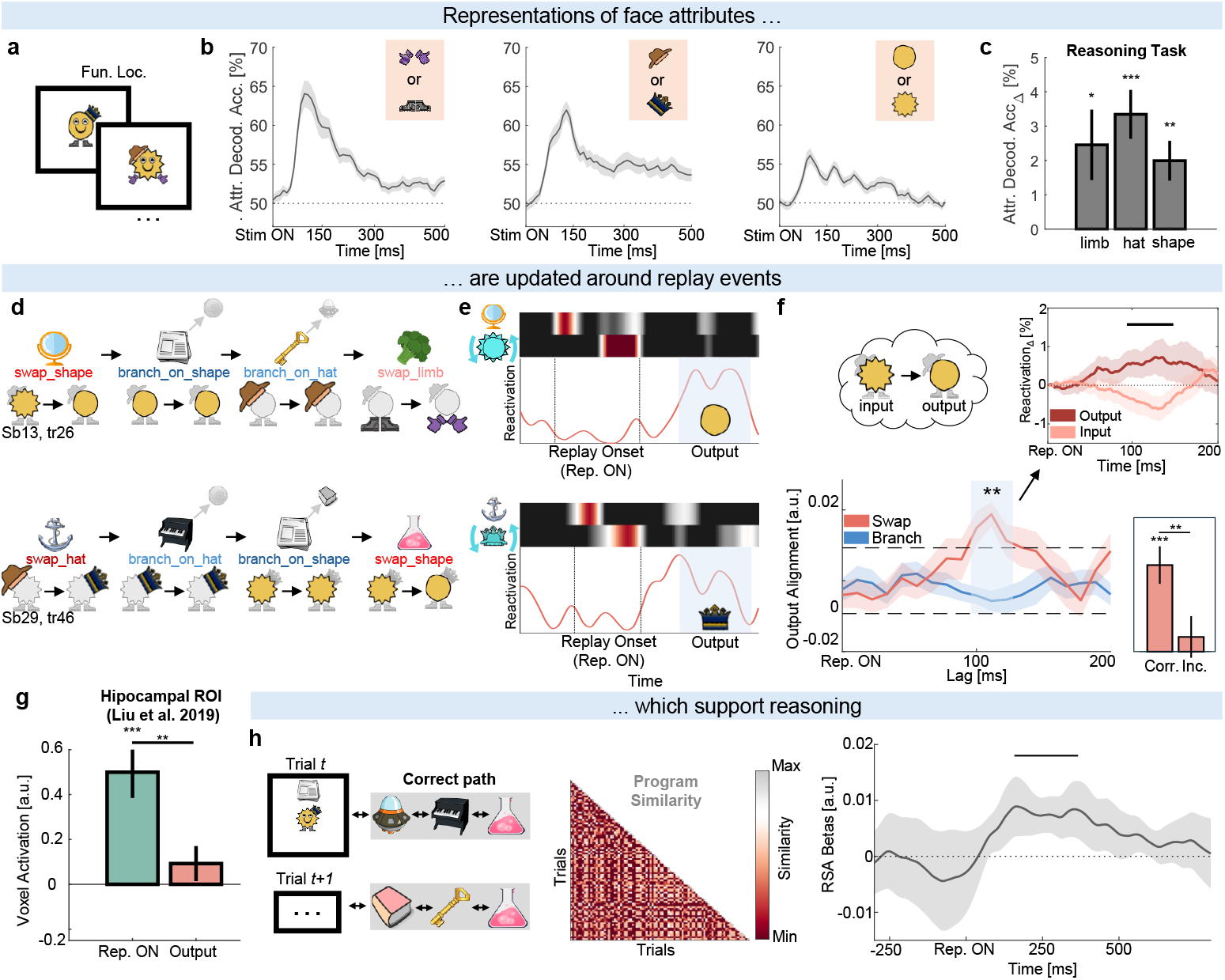
Replay sequences are followed by data updates and program similarity increases. ab) Face attribute decoding in the functional localizer. Second panel: binary classifier for [Hands, Feet]. Third panel: binary classifier for [Crown, Fedora]. Fourth panel: binary classifier for [Spiky, Round]. c) Peak face attribute decoding in the reasoning task at Reasoning Start for individual attributes. Time points used for averaging were determined using a leave-one-participant-out cross-validation procedure. d) Example program execution schematics for different trials and participants with highlighted data updates. e) Example Location → Operation sequences (“Replay Onset”) of correct paths that were followed by the predicted data update (Output). f) Location → Operation sequences containing swap operations (red shading) but not branch operations (blue shading) were followed by a data update. Dashed lines represent the 95th percentile of the null distribution (multiple-comparisons corrected). The bottom right figure inset shows the effect peak split for correct (Corr.) and incorrect (Inc.) trials. The top right figure inset shows input and output representations that were event-locked to the onset of replay events (see Methods: Program Execution with replay onsets). The black bar represents a significant difference between the output and input representation in cluster-based permutation testing (exceeding the 95th percentile of the mass-corrected null). g) Source reconstruction of ripple-band (80-180Hz) activity aligned to Location → Operation → Data sequences in the hippocampal ROI from Liu et al. ^1^. Sequences were split into Replay Onset (Location → Operation) and Output (Data). h) Left and middle panel: The program similarity regressor from Figure 4 was used to predict participants’ neural activity. The regressor was aligned to replay and control events (Supplementary Figure 5B). Right panel: Program similarity predicted participants’ neural activity after replay events (“Rep. ON”). The black bar represents significance in cluster-based permutation testing (exceeding the 95th percentile of the length-corrected null). *** p <.001, ** p <.01, * <.05. Mean values and thick lines represent the average across participants, shading and error bars represent the SEM across participants.

We used these decoders to ask whether Location → Operation sequences reported in the previous section were followed by data updates during reasoning. For example, would a Mirror → swap shape sequence (“Replay Onset”, Figure 7D, E) be followed by an update from Spiky into Round (“Output”, Figure 7D, E)? In this example, a systematic presence of Round would imply mental computation had been performed.

To look for evidence of Location → Operation → Data sequences we used TDLM whilst controlling for shorter sequences such as Location → Data or Operation → Data (see Methods: Program execution with TDLM). For this analysis to be meaningful, we grouped face attributes into Inputs (value before update; e.g. Spiky) and Outputs (value after update; e.g. Round). To measure the strength of the data update, we computed the difference between Location → Operation → Output and Location → Operation → Input sequences. This ensured operation and face attributes were always correctly matched and assumed participants correctly executed programs up to that point within a trial.

We found that Location → Operation sequences were followed by an update of the face attribute approximately 110-120 ms later (“Swap”, Figure 7F). This effect was primarily driven by shallow links, but was also significant for all links in the sequence (t_28_ = 3.39, p =.002, multiple-comparisons corrected). In line with correct program execution, it was present on correct trials (*t*_28_ = 5.41, p <.001, Figure 7F right inset) and appeared only for sequences involving *swap* operations. It was absent on sequences involving *branch* operations (“Branch”, Figure 7F): branches served as a control because they did not require a data update. Lastly, to provide further validity to this effect, we used a complementary approach where we did not use TDLM and instead event-locked neural activity to time points with significant Location → Operation sequences (see Methods: Program execution with replay onsets). We then computed reactivations of Input and Output attributes aligned to these events. This analysis corroborated our finding, revealing output attribute reactivations approximately 120 ms later (Figure 7F top right inset). While the results across both analyses are not independent (r =.70, p <.001), their convergence strengthens confidence in the effect.

Finally, we asked whether the observed sequences relate to hippocampal activity. Beamforming replay-aligned high-frequency (80–180 Hz, ripple-band) activity showed a significant activation increase in a hippocampal ROI previously associated with replay^1^ (*t*_28_ = 4.36, p <.001, Figure 7G). This activation was specifically observed during Location → Operation sequences (Replay Onset) but not during data updates (Output).

### 2.6 Program similarity strengthens around replay events

Previously, we observed that trials with similar programs acquired similar neural representations over the course of the reasoning period (cf. Figure 4C). We reasoned that participants mentally executed programs and stored the execution path in working memory. In line with our prediction that replaying sequences supports program execution, this effect was correlated with replay of program locations (r = -.43, p =.018, cf. Figure 5G). However, we were also able to conduct a more precise test of this idea: we asked whether each replay sequence is accompanied by a small increase in representational similarity between trials with similar programs.

We tested this by repeating the program similarity RSA analysis, but now event-locked to the onset of replay events (see Methods: Replay-aligned RSA analysis). A significant increase in program similarity was observed approximately 180-360 ms after a replay onset (Figure 7H) but not after onsets of control events (Supplementary Figure 5B). This pattern suggests - though does not prove causally - that replaying sequences actively supports program execution during reasoning.

## 3 Discussion

Reasoning involves composing pieces of knowledge to solve new problems. Recent theory has proposed that the brain composes pieces of knowledge by packaging them into fast spontaneous sequences^3^. To test predictions from this framework, we developed a task where participants executed short programs. Although the programs in our task were drawn from a restricted set, an appealing property of programs as a class of problem is that they can capture all possible formal reasoning^28^.

We found that by the end of execution, the similarity of inferred program paths predicted neural activity in prefrontal and parietal cortices, indicating participants correctly inferred and maintained the program solution in working memory. Importantly, the task associated locations in the program with visual objects. Throughout execution, object representations spontaneously reactivated in fast neural sequences. The patterns of reactivation were consistent with sequences sampling candidate partial solutions. Furthermore, spontaneous object reactivations were accompanied by abstract representations of operations at those locations, supporting the idea that replay sequences include information about the role of each item in the sequence. Lastly, internally generated sequences encoding a piece of the program were followed immediately by neural patterns reflecting the data generated by that piece of the program, hinting at an active role for sequences in executing programs.

During program execution, participants replayed program locations in patterns that align with previous human, AI, and rodent work^2;26;29–35^. For example, participants replayed program locations in sequences forming correct and incorrect paths. High-performing participants preferentially replayed correct paths, while low-accuracy participants replayed both correct and incorrect ones. When participants spent more time thinking, this was correlated with proportionally higher replay of incorrect paths. These results are in line with Schwartenbeck et al.^2^, where each replay sequence was a sample drawn from a space of possible combinatorial puzzle arrangements. They proposed a model where samples represent a mix of correct and incorrect combinations: when uncertainty is lower – such as in high-performing participants – the distribution of replay sequences tightens as samples become more confirmatory. This perspective also aligns with theoretical work where replay sequences represent samples of imagined actions^20^, and raises interesting questions about prioritizing samples^36^ in vast conceptual spaces in which they can be drawn arbitrarily without following paths on MDPs^2^.

Recent AI work^33–35^ shows models improve at sequential reasoning by backtracking from promising objects, tracking intermediate steps, and learning from productive mistakes (i.e. incorrect paths). Such strategies lead to more efficient reasoning through graphs because they promote learning internal world models^35^, in line with recent suggestions for hippocampal replay^4^.

We observed predominantly reverse replay. Like rodent tasks showing reverse replay^26;29–31^, our task was novel to participants, required working memory, and involved improvement throughout the session. Improvement implies ongoing learning about how to compose existing knowledge to solve the task. In rodents, reverse replay often reflects recently taken paths, likely to reinforce newly learned experience^32;36^. Related to these suggestions, participants replayed deeper program locations later in the session, which was mirrored in their behavior and consistent with Ou et al^37^ where participants preferentially replayed less known locations. Jointly, the similarities between our data and other human, AI, and rodent work suggest our task could be a rich testbed for bridging computational theories of planning and reasoning with their implementation in the brain.

Our RSA analyses suggest that the replay sequences we observed may have been part of a neural mechanism contributing to program execution. Furthermore, they suggest this process involves a network of brain regions (DLPFC, PPC, Crus II/I) involved in maintenance and manipulation of information in working memory^23;24^. The precise link between the patterns we observe and previously described reasoning mechanisms in cortex remains unclear, but our findings open several directions for future work^38–42^. Considering Location → Operation sequences were associated with increased activity in a hippocampal ROI previously linked to replay^1^, this raises the possibility that they reflect neuronal co-activations in hippocampus similar to those reported by Kunz et al.^43^. This leads to a broader question of whether hippocampal replay sequences could function as compositional “instruction sets” transmitted from medial temporal lobe to cortex (e.g. prefrontal cortex) where programs are executed^41^. In this view, cortical areas could integrate over time to decode the implication of the entities and roles assembled in a replay sequence, while also maintaining both the inputs and outputs of a replay sequence in working memory to perform computations complementary to hippocampal replay.

More broadly, our central findings contribute to a rapidly emerging understanding of the neural language of structured thought^44–49^. Binding of entities (semantics) and arbitrary roles (syntax) via replay allows the creation of structured representations that are a key ingredient for a language of thought^50;51^. The coexistence of rapidly composed structured representations and slowly learned cortical abstract representations implies a feedback loop where rapid composition in replay sequences allows discovering new knowledge that is used to train cortical abstractions^52;53^. In turn, these new cortical abstractions themselves may be used as inputs for higher-order compositional structured representations^49;54–56^. The presence of both factorized and compound sequences in our data would hint at this. Namely, such a loop may provide a mechanism by which the brain develops a generative grammar of increasingly complex representations, involving programs, that are used during reasoning. In summary, if we frame thought as the sequential reactivation of neural programs and our results generalize to other domains of cognition, we can begin to address fundamental questions about how we structure thought to reason.

## 4 Methods

### 4.1 Participants

Thirty-six healthy participants were recruited to participate in the study. All participants had normal or corrected to normal vision, had no known neurological or psychiatric illnesses and were fluent English speakers. Two participants were excluded after the first training session due to their performance being below a set threshold of 70%. The remaining 34 participants completed the study. Two participants were excluded from analyses due to artifacts in their neural data, likely due to dental metal, continuous large movements, or muscle tension. Three more participants were excluded from analyses due to their behavioral performance on the operation localizer being at chance, indicating they did not learn the required mappings between objects and operations. This left N = 29 participants (age range between 19 and 31 years old, 12 males) for analyses. Participants were recruited through UCL listings and a widely used participant pool (SONA) and were mostly undergraduate and graduate students. They all provided written informed consent prior to the start of the experiment and were financially compensated for their time (£10/h). Ethical approval was obtained from the Research Ethics Committee at UCL in compliance with national regulations, under ethics number 3090/004.

### 4.2 Functional Localizer Task

The functional localizer was used to train classifiers of objects and cartoon face attributes. The design of the functional localizer closely followed previous work^1^. On each trial, participants saw an image (object or face) for 0.8 seconds followed by a brief inter-trial interval period. On 10% trials, the image was followed by a text label to assess participants’ level of engagement. Participants had to provide an accept / reject answer whether the text label matched the previously seen image. The position of accept / reject was counterbalanced across participants. The text label was correct on 50% of trials. Images of objects were interleaved with images of faces in blocks of 50 trials. This was done to facilitate attention on objects vs. face attributes. In total, there were 38 trials per object presentation and 44 trials per face attribute presentation.

### 4.3 Reasoning Task

On each trial, participants were presented with a random pairing of starting location and face (STIM ON in Figure 2C). Based on the structure of the graph (Figure 2A) and associations between program locations and operations (Figure 2B) learned during training (see Procedure: Training), participants had up to 20 seconds to mentally execute the implied program until reaching the END token in their mind. Program execution was deterministic, which means that every program had one correct execution path (referred to as Correct Path). This meant, assuming participants correctly executed programs, we could probe the exact sequence of program locations, their operations, and face attribute updates on every trial. Participants were able to terminate the reasoning period early via button press if they thought they reached the END in their mind. Unbeknownst to the participants, there was a lower bound of 5 seconds that was used to ensure participants would not rapidly terminate trials due to lack of motivation. After the reasoning period was over, participants were asked two probe questions: END and PATH questions. END questions assessed participants’ knowledge at the END token, providing us with information whether their knowledge of the last “encountered” object or face was congruent with the predictions of the correct path. Similarly, PATH questions provided us with information whether their knowledge of the objects and faces “encountered” on their execution path were congruent with predictions of the correct path. Across trials, both question types (END / PATH) were randomized in presentation order, in addition to being randomized and equally balanced in question modality (object, face). Within trials, question modality was kept constant across END / PATH questions to minimize participant confusion. Participants had 3.5 seconds per question to provide an accept / reject response with 50% trials showing the correct image (i.e. object or face) or an incorrect one. Participants received no feedback about the accuracy of their response. In total, there were 88-92 trials per participant (across 8 blocks) with 42 unique sampled correct paths on average across participants. Here, unique correct paths strictly refer to the number of unique paths stemming from the starting combination of program location and face attributes. This highlights the novelty of individual trials throughout the task. Unbeknownst to the participants, correct paths would vary in length, requiring participants to reason through sequences of three or four objects / operations. Length was varied to prevent over-learning of possible paths and ensure participants did not consider the length of a program a vital component of the task. Accept / reject button bindings were counterbalanced across participants.

### 4.4 Operation Localizer Task

The operation localizer was used for testing how well participants memorized the association between objects and operations and for training classifiers of their corresponding abstract representations: operation type (swap or branch), operation attribute (“limb”, “hat”, “shape”), and the specific operations (swap limb type, swap hat, swap shape, branch on limb type, branch on hat and branch on shape). The structure of the operation localizer was similar to the functional localizer and closely followed previous work^1^. Participants saw an object for 1.4 seconds and were required to think about its operation. Afterward, they were required to accept / reject the indicated operation (text label) that appeared on the screen for 2 seconds. Participants had to provide an accept / reject response whether the label matched the previously seen object. In 60% of the cases, the operation text label matched the object. The slight positive bias was introduced to reduce the probability of participants becoming uncertain about the mapping between object and operations and thus reducing the fidelity of these neural signals. Each object was presented 22 times, yielding 44 presentations per operation.

### 4.5 Procedure

The whole study lasted two training sessions (Training: Day 1 & Day 2) and one testing session (Testing: Day 3), carried out in the MEG scanner. During two consecutive days of training, participants were taught the building blocks to solve the reasoning task. The experiment was framed as a memory-based card game in which they would first need to learn a deck of cards and their abilities over two consecutive days before playing the game in the MEG scanner on the third day.

### 4.5.1 Training: Day 1 & Day 2

Due to the high complexity of Day 3, we used different subtasks on Day 1 & Day 2 to teach participants all the necessary components. For participants to successfully reason on Day 3, they were required to combine their knowledge from both subtasks in continuously novel ways.

On Day 1, participants were taught transitions between 12 objects and one END token. A session lasted approximately 60 minutes and was largely self-paced. This involved showing participants an object (depicted by an image) and its links (e.g. Newspaper → Spaceship or Spaceship → Key or Piano). Participants would subsequently see an object (e.g. Newspaper) and had to indicate its correct link from two possible objects (e.g. Spaceship and Broccoli). The same pairwise transition logic was applied for all objects. The goal of this training procedure was to ensure participants understood pairwise transitions between objects. Towards the end of Day 1, participants were shown longer sequences of objects that varied in length (n = 108 trials). Participants were required to indicate whether a sequence was correct or not. The goal of this training procedure was to ensure participants understood that objects with two possible links could not form a correct sequence with both links in the same sequence. This was important because six objects transitioned to one further object (or token) and six objects transitioned to two further objects (or one object and one token). For example, if the possible transitions from Spaceship were Spaceship → Piano and Spaceship → Key, then Spaceship → Piano → Key or Spaceship → Key → Piano were not possible correct sequences. Instead, a valid sequence could only be Spaceship → Key → … or Spaceship → Piano → …. Feedback about the objects comprising the sequence was provided if participants answered incorrectly. This was done to facilitate learning. By the end of Day 1, participants learned there were 12 objects in total and what their transition structure was. Some objects connected with a terminal END token. They were not informed about the logic behind END nor the reason objects could transition to a different number of objects.

On day 2, participants were taught the associations between objects and operations. Operations defined the computations participants had to execute when encountering an object. Participants were further taught about the existence of a data structure visually represented by a cartoon face. Each face was a composition of three orthogonal attributes (limb: hands or feet, hat: fedora or crown, shape: round or spiky) each depicted by an image fragment. In total, there were eight unique faces which served as “input” data for the operations participants executed. For example, on Day 2 participants learned objects were associated with “swap” operations. This operation required them to swap one of the three attributes of the face. If they would see [spiky, hands, crown], “swap hat” required updating crown to fedora and vice versa. The same logic applied to all other attributes. Participants also learned objects were associated with “branch” operations. Learning about swap and branch operations contextualized why some objects transitioned to two objects while others transitioned to one. For example, on Day 1 a participant might have learned Spaceship → Piano and Spaceship → Key are valid transitions. On Day 2, they might have learned Spaceship required them to “branch on hat”. The transition from Spaceship that would be correct *on a given trial* depended on the current value of the “hat” attribute. Thus, for objects with branch operations, participants effectively learned “if-else” conditionals: encountering Spaceship and [spiky, hands, crown] would lead to Key, while [spiky, hands, fedora] would lead to Piano.

The goal of Day 2 was learning the associations between objects and their operations, and how these interacted with face attributes. These operations were consistent with the graph learned on Day 1 - each swap object transitioned to one object and each branch object transitioned to two objects. The structure of the training was similar to Day 1 with the addition of operations: participants saw a combination of object and face and had to indicate the correct combination of object and face that followed. On Day 2, they did not see passive sequences like on Day 1 to avoid contamination and retain novelty on Day 3. Instead, participants learned one-step transitions as this was the core reasoning mechanic they would have to chain together on Day 3. At the end of Day 2, participants were instructed on the structure of the reasoning task they would do while in the MEG scanner on Day 3. They were given a few trials for familiarization with button bindings and its logic. Those trials were not used on Day 3 to ensure trial novelty while participants were in the MEG scanner.

Across participants, the positions of objects and their operations were randomized. We intentionally used objects and attributes with low semantic similarity (Supplementary Figure 1A) to minimize biased graph learning or spontaneous generation of associations between objects / attributes due to high semantic similarity. Across participants, we counterbalanced four possible graph structures (Supplementary Figure 1B).

### 4.5.2 Testing: Day 3

On Day 3, participants could refresh their knowledge of the graph structure for 5-10 minutes by seeing individual objects, their operation associations, and objects they transitioned to. They were then carefully instructed about the whole procedure of Day 3. Our experimental design on Day 3 was conceptually analogous to Liu et al.^1^: participants first underwent the functional localizer, followed by the reasoning task, and finished with the operation localizer. After the experiment, we debriefed participants and asked them about the strategies they used to solve the task, the strategies they used to learn the graph structure, and probed their knowledge of the graph structure. The whole session on Day 3 lasted approximately 2 hours and 30 minutes, 2 hours of scan and preparation times, in addition to 30 minutes for the instructions and debriefing after the scan.

### 4.6 Behavioral analyses

Participants’ behavior on all Day 3 tasks (functional localizer, reasoning task, operation localizer) were analyzed. For the reasoning task, we computed participants’ accuracy as the probability of providing a correct answer (i.e. correctly accepting or correctly rejecting) to the object or face probe appearing during END / PATH questions. Depending on the behavioral analysis, trials were split based on the key condition of interest (question type, question modality, program length, and their interaction). We also examined participants’ thinking duration as a function of program length and the number of branches occurring within the program. To compute the interactions between number of branches (or swaps) on accuracy and thinking time (Figure 3G-I and Supplementary Figure 2B), we first computed these quantities across trials of identical program length and then averaged across program lengths. This ensured the effect was not confounded by accuracy decreasing or thinking time increasing on longer programs.

In Supplementary Figures 2 and 3, we computed the effect of attribute distance on END Face questions and the effect of depth on PATH Object questions, respectively. To compute the latter, we computed the probability of correctly accepting the probe questions when they were sampled from the correct path. We split them depending on the location in the correct path from which they were sampled (Shallow or Deep). To compute the former, we split trials based on the distance of the sampled probe from the correct face. That is, “0” corresponded to trials where the sampled face probe matched the correct END face and thus required participants to correctly accept. Other values corresponded to trials where sampled face probes were one to three attributes different from correct END faces, thus requiring rejection. In the functional and operation localizer, participants’ accuracy was defined in an identical manner to the reasoning task (accept / reject).

### 4.7 MEG acquisition and preprocessing

MEG recording took place at the Wellcome Centre for Human Neuroimaging using a whole-head 275-channel axial gradiometer system (CTF Omega, VSM MedTech). The data were acquired continuously at a sampling rate of 600 Hz. Eye movements were also simultaneously recorded using an eye tracker to help identify artifacts in preprocessing. Prior to the MEG session, participants were given detailed instructions about the experimental task and were screened for any contraindications such as metallic implants. Participants were seated upright and reminded to stay as still as possible. They made responses using their right hand on a four-by-one button box. Each participant completed 14 experimental blocks, three functional localizer blocks, followed by eight reasoning task blocks, followed by three operation localizer blocks. A full MEG scan session lasted between 100-120 minutes, including break times and set-up time. The preprocessing protocol closely followed previous work^1;2;22^. The data were downsampled from 600 to 500 Hz (reasoning task) and 100 Hz (functional and operation localizer) to optimize processing time and signal-to-noise ratio. All data were then high-pass filtered at 0.5 Hz using a first-order IIR filter to remove slow drift. Artifacts caused by rapid eye movement, muscle clenching, and excessive head movements were then identified through visual inspection and automated pipeline trained to recognize such patterns. These segments were labelled as ‘bad’ and excluded from the analysis by subtracting them from the data to be analysed. An independent component analysis (ICA) was used to decompose the sensor data for each session into 150 temporally independent components and associated sensor topographies. Further analyses were conducted on the preprocessed MEG signal measured in units of femtotesla and collected across sensors covering the entire brain.

### 4.8 Neural analyses

#### 4.8.1 RSA analyses

We investigated whether the similarity of correct program paths across trials (“Program Similarity”) predicted participants’ neural activity using a GLM-based representational similarity analysis (RSA). Program similarity measured the position-dependent overlap of objects between trials. For example, the paths 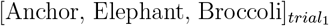 and 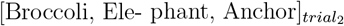 would have low program similarity because only Elephant is in the same position.

Note that program similarity was only computed using program locations that were *not* visible to participants during the reasoning period. That is, the starting location - Newspaper in Figure 4A for example - was not included. Thus, across-trial program similarity could only predict neural activity if participants inferred and mentally navigated through the graph during the reasoning period.

The GLM containing the program similarity regressor additionally included a bias term and regressors controlling for similarity arising from the correct face path and differences in program length. The latter was used to account for the fact that differences in similarity could also arise due to differences in length. When program similarity between trials of different length was compared, we assessed similarity separately for the first object, middle objects, and final object.

In addition to the above GLM, we conducted a further analysis in which we added a regressor that measured the position-independent overlap of objects between trials (“Object Similarity”). While program similarity for [Anchor, Elephant, Broccoli]_*trial*1_ and [Broccoli, Elephant, Anchor]_*trial*2_ is low, object similarity is high because all three objects appear on both trials. We used both the program and object similarity regressors in the same GLM in Figure 4D.

To distinguish contributions of Shallow and Deep program locations to the program similarity effects observed in Figure 4, we split each trial’s correct path based on the association of the starting location (Shallow) and associations of Shallow (Deep). For example, if a trial started with Newspaper, as in Figure 4A, Shallow measured similarity based on its association (Spaceship), while Deep measured similarity based on objects after Spaceship (Piano, Beaker).

The empirical neural similarity matrix was obtained by computing the across-trial correlation between MEG sensors at each time point on smoothed neural activity^25^. Because trials were self-paced, we analysed two key epochs: the reasoning period start (STIM ON; Figure 2C) and end (STIM OFF; Figure 2D). Reasoning Start was aligned to the onset of each trial - when the starting location and face were first presented. In contrast, Reasoning End was aligned to the offset of each trial - when both stimuli were removed from the screen and replaced with a fixation cross before participants were probed with END / PATH questions. For each period and participant, we obtained a time series of RSA beta coefficients. Statistical significance of these beta coefficients was determined by examining whether they form clusters exceeding the 97.5th percentile of a mass-corrected null distribution (n = 1000 shuffles) which was obtained from random shuffles of neural data before recomputing the RSA betas.

#### 4.8.2 RSA Source reconstruction

To localize the putative anatomical sources of our effects, we followed a similar approach to previous work^1;22^. Sensor-level data were projected to a three-dimensional grid in MNI space (grid step, 5mm) using linearly constrained minimum variance beamforming as implemented in OSL^57^. Forward models were computed on the basis of a single shell head model using superposition of basis functions that approximately corresponded to the plane tangential to the MEG sensor array. The sensor covariance matrix was estimated on the cluster significant in permutation testing (Figure 4C) around the Reasoning End period using the smoothed wideband (1-180Hz) MEG data. At the first level, we computed one-sample tests on wholebrain source activity. For group-level inference, participant-level t-maps were smoothed in OSL using a 5-mm FWHM Gaussian kernel and entered in a one-sample t-test to estimate the strength of a main effect compared to a 50 ms baseline before the significant cluster^1^. We then performed whole-brain cluster-level correction and visualized clusters significant at (p <. 05).

### 4.8.3 Functional localizer decoding

To look for spontaneous reactivations of objects and face attributes, we trained binomial classifiers, closely following the approach from previous work^1;21;22;25^. The functional localizer data were split into trials with object and face presentations. On the two different trial splits, separate binomial classifiers were trained using a leave-one-out approach for each object and face attribute. The binomial classifier for objects was trained such that all trials except one for each object were used as a positive training example, all other objects as a negative training example. One object trial was held out as test data. This was repeated for all objects (Chance decoding: 8.3%). A similar approach was used for face attributes where binomial classifiers were created for each face attribute. In this case, pairwise classifiers were used (i.e. all trials of ‘hands’ except one were used as a positive training example and all trials of ‘feet’ were used as a negative training example), yielding chance decoding at 50%. This approach was repeated for all objects and all face attributes. The decoding accuracy – the probability of decoding the correct object / face attribute on held-out data – was computed for each participant, smoothed, and averaged across participants. A confusion matrix was then constructed from the peak decoding window across participants for object decoding accuracy. The peak decoding windows observed in our data are comparable to those reported in previous similar work^21;22^. Decoding accuracy significance was assessed via mass-corrected permutation-testing. The true object label was shuffled (n = 1000) across trials for each participant to build a null distribution. Significance was determined by examining whether the empirical decoding accuracy across participants forms clusters exceeding the 97.5th percentile of the mass-corrected null distribution.

#### 4.8.4 Sequenceness analyses

The strength of sequential reactivations was measured using temporally delayed linear modelling (TDLM), as in previous work^1;2;19;25;58^. Using the trained classifier weights, we computed reactivation probabilities for objects, operations, and face attributes during the reasoning period of the reasoning task. To compute sequenceness, we applied a two-stage GLM approach. At the first stage, we generated empirical transition matrices between the representations of interest (e.g. objects, operations, face attributes) on a trial-by-trial basis. Because transition matrices were always defined on a trial-by-trial basis, the sequenceness estimates obtained from TDLM analyses in our data were particularly susceptible to artifacts remaining after preprocessing. Therefore, as an additional preprocessing step, we calculated the entropy of reactivation probabilities and removed trials if an individual object was almost never reactivated during a trial’s reasoning period (approximating a reactivation frequency between 2.5 and 5% of all time points). Across participants, this preprocessing step removed 1.04% trials on average. We computed the empirical transition matrices, measuring the strength of sequential reactivations, using different time lags up to 200 ms based on the effects observed in previous work^1;2;22;25;37;58^ and averaged coefficients of neighboring lags to further improve statistical power. At the second stage, we then aimed to predict the empirical transition matrices using theoretical transition matrices while controlling for self-transitions in addition to using a bias term. Significance was computed by creating a null distribution by shuffling (n = 1000) the state identity of the theoretical transition matrix of interest. This had to be repeated for each trial and for each participant because theoretical transition matrices changed on a trial-by-trial basis. For each participant, we averaged the obtained sequenceness values across trials and then across participants. We computed the 95th percentile of the null distribution from these averages and used the maximum value across lags to correct for the numerous tested lags.

#### 4.8.5 Location → Location sequences

In Figure 5G-I, correct paths were defined as the correct execution path within a 12x12 object transition matrix, specified on a trial-by-trial basis for each participant. Using the example from Figure 5D, while the correct path on trial 1 would have comprised [Newspaper, Spaceship, Piano, Chemistry], for example, this would have been different on trial 2 (e.g. [Book, Anchor, Elephant, Broccoli]). Because we shuffled object identity to create the null distribution, trial-wise transition matrices explaining participants’ neural activity would imply participants replayed program locations in sequences adhering to paths on the graph during reasoning.

To analyze the contribution of individual locations to the correct path sequenceness, we split the correct path into its constituent links (Shallow, Deep), like in *RSA analyses*. We then computed the strength of sequenceness at the peak lags (70-90msec) and did a t-test across participants. To make sequenceness estimates comparable across participants and trials, we z-scored them within participant by their respective null distribution. To compare the patterns in participants’ replay sequences, we split participants (median split) based on their overall accuracy on the probe questions.

Incorrect paths were defined as all transitions that were possible between objects given the 12x12 transition matrix structure of the object graph on trial *t* (excluding the correct path on that trial). For example, on Trial 1 these would involve [Mirror, Book, Key, Elephant, etc.]. Note that an incorrect path on one trial could be a correct path on another trial and vice versa.

### 4.8.6 Independent dataset analysis

We used data from Schwartenbeck et al.^2^ to test whether we could replicate the effect reported in Figure 5I in an independent dataset. Briefly, in Schwartenbeck et al.^2^ participants (n = 20) were required to solve combinatorial puzzles. Each puzzle was composed of different building blocks that could be a) *Stable* (appearing in all puzzles), b) *Present* (and connecting to Stable), c) *Present* (and not connecting to Stable), or d) absent (the building block did not appear in a particular puzzle). Their work showed that while participants’ replay sequences initially tested out various possible solutions to the puzzles they had to solve (early time interval in Figure 7B from Schwartenbeck et al.^2^), the sequences eventually converged on the correct solution (late time interval in Figure 7B from Schwartenbeck et al.^2^). We reasoned that if across both datasets we are measuring a similar neural phenomenon, we could conceptually replicate our finding from Figure 5I despite differences in tasks. Therefore, we repeated an analogue of our analysis by splitting the sequences reported in their data into “Correct” and “Incorrect” samples. This was done on the late time interval where only correct samples were reported as significant across participants. We then split participants based on their accuracy in solving these puzzles (median split) and examined whether their main claim was related to participant accuracy, like in our data. Lastly, the sequenceness values obtained from their data were multiplied by -1 to ensure conceptual similarity to our data. Due to the design, in their data the direction of the effect was arbitrary. That is, forward *Present* → *Stable* sequences could also be interpreted as backward sequences between the building block that was always visible (*Stable*) and the candidate building blocks that it could connect to (*Present*). In our data, this would be conceptually analogous to observing a backward sequence from Piano to Newspaper when Newspaper was the starting location, for example.

#### 4.8.7 Operation localizer decoding

To test whether participants developed abstract representations of operations, we trained binomial classifiers using a generalized version of the approach from Liu et al.^1^, where abstract codes such as “position” or “graph index” were observed. The central idea underlying this approach is that if two objects (e.g., Newspaper and Broccoli) share an abstract representation, then a classifier trained on Newspaper trials should decode above chance on Broccoli trials. Because object–operation assignments were randomized per participant, any successful generalization across objects could only reflect abstract operation codes rather than object-specific visual features.

We built three neural classifiers. The first targeted operation type (swap vs. branch), requiring a shared neural representation of “swap” and “branch” that generalizes across both objects and attributes. For example, training involved using trials containing branch on hat and branch on shape as positive examples, while swap hat and swap shape trials were used as negatives. The classifier was then tested on held-out trials involving the remaining attribute (e.g., branch on limb type vs. swap limb type). Because the train/test split did not involve shared attributes, we were also able to build a classifier targeting operation attribute (hat / limb / shape). For example, branch on hat trials were used as positive examples, while branch on shape and branch on limb trials were negatives. Testing was performed on swap hat trials as positive examples. Above-chance decoding here would indicate an attribute-specific “hat” representation that generalizes across both branch and swap operations. We also trained classifiers for specific operations that combined both representations (Type and Attribute). For each specific operation (e.g., swap hat), one object associated with the operation was used for training and another object associated with the same operation was used for testing, requiring generalization of the specific operation across both objects. To illustrate using the example from Figure 6CD, training for operation type involved using trials with [Piano, Coin, Broccoli, Newspaper] and testing on trials with [Web, Anchor]; training for operation attribute involved using trials with [Web, Anchor] and testing on trials with [Key, Mirror]. Lastly, training for specific operations involved training on trials with [Piano] and testing on trials with [Coin]. Across all analyses, train/test splits were repeated combinatorially to exhaust all possible partitions. Because the data used for training each of the described classifiers was different, the indexed representations were statistically independent.

Decoding accuracy significance was assessed in an identical manner to the one described for *Functional Localizer Decoding*.

#### 4.8.8 Abstract representation decoding

To validate that classifiers trained on the operation localizer could reliably decode abstract representations that generalize to other task epochs, we used data from the functional localizer and the reasoning task. Because the functional and operation localizer structure was identical (a single centrally presented image), we computed decoding accuracy over time and assessed significance as in *Functional Localizer Decoding*.

To evaluate decoding accuracy in the reasoning task, however, we had to account for across-task differences in the design. First, the starting location and face were presented simultaneously at reasoning start. Second, the face was presented centrally while the object was presented at the top of the screen. The object’s placement differed from the localizers’ central presentation and required saccading for overt attention. To account for this, we temporally aligned neural activity to participants’ saccade patterns before computing operation decoding accuracy. If no saccade information was available, the across-trial median was used for temporal alignment. In contrast, while only one centrally presented image was shown to participants at the end of the reasoning period (END / PATH questions), they could have formed predictions about which END / PATH object may appear based on having executed programs during the reasoning period. Thus, to be able to meaningfully average across both reasoning task epochs to maximize statistical power and avoid false positives, we estimated decoding accuracy peaks for the Type and Attribute classifiers using a leaveone-participant-out cross-validation approach. We estimated the peaks on n-1 participants, stored the peak information and used it to extract the decoding accuracies of the held-out participant. Based on Liu et al.^1^ and the temporal profile of these representations in our localizers, we used a 300 ms window at both epochs to estimate the decoding accuracy peaks. This procedure was repeated for both epochs across all participants. In Figure 6G, the across-epoch averages are presented.

#### 4.8.9 Validation simulations

Noting that the operation decoders consistently reached decoding performance 1-2% above chance in cross-validation (Figure 6EFG), we conducted simulations to confirm that this level of accuracy is sufficient to decode operations in replay.

We created synthetic localizer and reasoning neural data of n = 20 simulated participants. All synthetic data used temporally autocorrelated Gaussian noise designed to approximate real neural data in terms of comparable trial durations, trial numbers, and sensor numbers. On each trial of the localizer and reasoning data, we added discrimination patterns on 10% of sensors that were used to encode the operation presented on that trial. In simulated reasoning data reactivation matrices, we then added Location → Operation sequences at random times. Within each trial, a simulated sequence consisted of object reactivations (n = 50) followed by operation reactivations at lag *X* where *X* ∼ 𝒩 (30, 10) ms. The operation representations were defined as the same patterns over sensors used in the simulated localizers.

We then applied an identical analysis pipeline to the simulated data. First, we measured accuracy in decoding operations in cross-validation on the localizer. Accuracy reached 2.01% above chance level (Supplementary Figure 4C). Next, we measured Location → Operation sequenceness in the synthetic reasoning data reactivation matrices. We found a strong peak in sequenceness which exceeded the multiple comparisons-corrected permutation threshold (Supplementary Figure 4D green line). As a control, we also analyzed reasoning data reactivation matrices where no real sequences were added, and confirmed that the analysis did not produce false positives (yellow line). We conclude that in simulated data, decoders with comparable levels of accuracy in cross-validation are sufficiently strong to decode such sequences.

#### 4.8.10 Location → Operation sequences

Analyses in Figure 6 examining whether reactivations of program locations were accompanied by operations followed the same procedure as described in *Sequenceness analyses*. The key difference here is that the theoretical transition matrix included transitions between objects and operations rather than between objects themselves. The theoretical transition matrices therefore included reactivations of both objects and operations. These were defined by the factorized (Type, Attribute) representations which we tested in separate GLMs (first two analyses), together in the same GLM (third analysis), and by the specific operation representations themselves (fourth analysis). Importantly, classifier weights used to generate operation reactivations were independent of the object image: for example, if a trial contained Coin (swap limb), we used classifier weights from Piano (swap limb) to compute reactivation probabilities. This train/test logic was applied in operation-related analyses to avoid false positives related to operation reactivations that could arise from decoding of an object’s identity and or visual features.

Operation-related reactivations were always conditioned on the correct object path, like in *Location Location* → *sequences*. Note that because classifier weights were trained on independent data, both bars in Figure 6K, the result in panel M, and the result in panel S represent three independent tests. Furthermore, note that because Type and Attribute comprised only two and three features respectively, we restricted these analyses only to correct trials in which the path did not repeat a feature combination to ensure no contamination by visual features (see *Operation Localizer Decoding*). Using the empirically observed lags from Location → Type and Location → Attribute sequences, we could specify a single design matrix in which we combined the regressors of each dimension in one GLM ([Location → Type → Attribute]). This meant we looked for reactivations of factorized sequences in close analogy to those observed in previous work^1^.

While statistical significance across the first three analyses was determined using the same procedure as in *Sequenceness analyses*, we used additional cross-validation approaches to determine the robustness of our results. For the first two analyses, we used a leave-one-participant-out cross-validation to determine the significant Location → Type and Location → Attribute lags on n-1 participants. We then used those lags to extract the sequenceness estimates on the held-out participant and repeated this procedure for all participants. In Figure 6K, we report the results of the cross-validated held-out sequenceness estimates (left panel) and the distribution of lags across all cross-validation folds (right panel). Because factorized sequences reported in the third analysis (Figure 6M) are predicted by the effects from the first two analyses (Figure 6K), we also used a cross-validation procedure where we estimated the effects from the first two analyses on one half of the data and asked whether they predict the lag structure of the third analysis on the other half of the data. Specifically, we estimated Location → Type and Location → Attribute sequences on even trials, and asked whether they predicted the lag structure of the factorized sequence on odd trials. We then repeated the procedure in the opposite direction (odd trials for the first two analyses, even trials for the third analysis).

In Supplementary Figure 4F, we conducted another analysis to provide corroborative evidence for the representational structure reported in Figure 6J-N. Because replay can be thought of as time-compressed reinstatement of memory content, we extracted the peak decoding latency for object, Type, and Attribute representations for each participant. We then calculated the percentage of participants whose peak latencies followed the predicted Location → Type → Attribute order as reported in Figure 6J-N. This empirical proportion was compared to the 97.5th percentile of a null distribution where the label identity (object, Type, Attribute) was randomly shuffled (n = 1000) within participant before computing the representation order pattern.

Because the specific operations in the fourth analysis can be viewed as conjunctions of the Type and Attribute dimensions, we asked whether the effect of the fourth analysis strengthened with experience and ongoing consolidation during the task. To test this, we quantified how the Location → Operation sequence effect evolved across the session. Specifically, we computed the effect within a sliding window, averaging over the first third of trials, incrementing the window by one trial, and repeating until the end of the session. We then fit a trial-index regressor to these values, yielding a beta coefficient that captured the slope of change across the session. Positive beta coefficients indicate increasing sequence strength, whereas negative coefficients indicate decreasing strength.

#### 4.8.11 Program execution with TDLM

To measure whether attributes of the face data structure were updated following Location → Operation sequences, we employed multi-step TDLM^1;2;58^. The procedure was identical to one described in *Sequenceness analyses*, with the addition of explicit controls for shorter sequences. In brief, we tested whether Location → Operation sequences were followed by updates in the face attribute representation predicted by the operation. For example, a Mirror → swap shape sequence should be followed by a Spiky → Round update. Explicit controls ensured that effects were not driven by shorter Location → Attribute or Operation → Attribute sequences. Specifically, we controlled for sequences where object reactivations were directly followed by attribute reactivations (omitting operation reactivations), and for sequences where object reactivations were followed by attribute reactivations at lags consistent with a missing operation reactivation.

This analysis fixed the Location → Operation lag based on results from Figure 6 and tested whether an update of the face attribute representation occurred in the subsequent 200 ms. The update was assessed by comparing sequenceness strength for Location → Operation → Spiky (input) and Location → Operation → Round (output). To obtain a net measure of representational updating, these values were subtracted (Output – Input) and averaged across trials within participant. Significance was evaluated by shuffling input/output labels within participant (n = 1000) and applying the same multiple-comparison correction as in *Sequenceness analyses*.

Because participants’ PATH Face and END Face accuracy decreased as the number of swaps in a sequence increased (Figure 3I, Supplementary Figure 2B), we separately examined the strength of this effect across correct trials of all links, and shallow links specifically (Figure 7F).

Due to our task design, we were able to extend this analysis to a control condition containing branch sequences. This allowed testing whether the predicted data update was specific to sequences containing swap operations (where an update was expected) or whether it would also occur on sequences containing branch operations (where no update was expected). Like above, inputs were defined as face attribute values before a hypothetical data update and outputs as the same values after a hypothetical update.

#### 4.8.12 Program Execution with replay onsets

To define replay onsets, we closely followed methods from previous similar work^1;22;25^. On each trial, we first identified time points with significant Location → Operation sequences. We did this by multiplying the object reactivation at time *t* (*X*_*t*_) with a corresponding time-lagged operation reactivation (*X_δt_*) where *δ* represented the empirically observed lag from Figure 6. This was done on all time points of each trial of the reasoning period for every object comprising the correct path for each participant. To determine significant reactivations, identities of all elements (objects/operations) were shuffled (n = 1000 shuffles) before multiplication to generate a null distribution. Replay onsets were identified as events where the empirical joint reactivation probability surpassed the maximum value of the reasoning period’s 95th percentile of the null distribution. Finally, we excluded replay events where another replay event occurred within a 200 ms window to avoid overlap.

These replay events (“Replay Onset” in Figure 7) were used to provide corroborative evidence for program execution. This was done by temporally aligning and epoching input and output reactivations to each event, smoothing, and baseline-subtracting to ensure a common baseline across events, trials, and participants. Events were then averaged within trials. Finally, across-participant averages in the input and output representation time courses were compared. Specifically, because this analysis is conceptually similar to Program execution with TDLM (without the controls for shorter sequences described in the previous section), we were interested whether an increase in the output reactivation would temporally coincide with the “Result” effect (Figure 7F). To test this, we investigated if the difference between the output and input representation was significantly stronger compared to chance by shuffling (n = 1000) the label (input / output) and determining if a cluster exceeding the 95th percentile of a mass-corrected null distribution could be detected.

#### 4.8.13 Replay-aligned RSA analysis

Evidence for a relationship between replay onsets and program execution was also obtained using a replay-aligned RSA. Specifically, we tested whether neural activity following replay events was explained by the program similarity regressor from Figure 4, as described in *RSA analyses*. To do this, we temporally aligned participants’ neural activity to replay onsets described above. Because multiple replay events could occur within a trial, we first smoothed and averaged neural activity across all replay events within each trial, computed separately for each link. This then allowed conducting the same RSA analysis described in *RSA Analyses* and examining the time course of RSA betas aligned to replay events.

Statistical significance was assessed by shuffling neural data (n = 1000) prior to RSA estimation and testing whether clusters exceeded the 95th percentile of the length-corrected null distribution. As a control, we repeated the analysis using “control events,” defined by randomly sampling time points during the reasoning period within each participant. The number of control events matched the number of replay events and could overlap with replay periods. Because control events were sampled randomly, we repeated the full process for each control onset within each trial of each participant - several times to maximize robustness (n = 5; Supplementary Figure 5B).

#### 4.8.14 Input representation decoding

In the reasoning task, the classifiers trained on face attributes were used to compute the decoding accuracy for the relevant attribute at the beginning of each trial (Reasoning Start). The “input” representation in Supplementary Figure 5A always referred to the attribute that the operation of the starting location operated over. For example, if a trial started with Spaceship (Branch Start), participants needed to maintain the hat-related representation (e.g. Crown) to be able to conditionally branch to the next program location. In contrast, if a trial started with Newspaper (Swap Start), they had to update the input (e.g. Crown) to an output (e.g. Fedora). To test if the input representation was maintained on Branch Start trials, we compared the decoding accuracy to a null distribution where the input / output [e.g. Crown / Fedora] label was shuffled (n = 1000) across trials per participant to investigate if we would detect clusters exceeding the 97.5th percentile of a mass-corrected null distribution. To test if the input representation was significantly stronger on Branch Start vs. Swap Start trials – indicating participants were sensitive to the requirement of the starting location’s operation - the trial label (Branch / Swap) was shuffled (n = 1000) and the difference between them computed. We used an identical mass-corrected permutation-testing approach as mentioned above to determine statistical significance.

## Acknowledgements

We thank J. Selvanayagam, C. Lindersson, M. El-Gaby, and A. Mahmoodi for very helpful discussions. We thank Chloe, Dan, Dimitra & Yasmin for technical assistance with MEG recordings and the volunteers who took part in the study. SV was supported by the Leverhulme Doctoral Training Programme for the Ecological Study of the Brain 445 (DS-2017-026). EG was supported by an MRC grant (MR/N013867/1). TEJB was supported by a Wellcome Trust Senior Research Fellowship (104765/Z/14/Z), a Wellcome Trust Principal Research Fellowship (219525/Z/19/Z), a James S. McDonnell Foundation grant (JSMF220020372), a Wellcome Trust Collaborator award (214314/Z/18/Z), the Gatsby Initiative for Brain 450 Development and Psychiatry (GAT3955), and by the Jean Francois and Marie-Laure de Clermont Tonerre Foundation. SWK was supported by the National Institute of Mental Health (F32MH081521), Wellcome Trust Investigator Awards (096689/Z/11/Z, 220296/Z/20/Z), and a BBSRC Strategic Longer and Larger Grant (BB/W003392/1).

## 5 Supplementary Figures

**Supplementary Figure 1:**
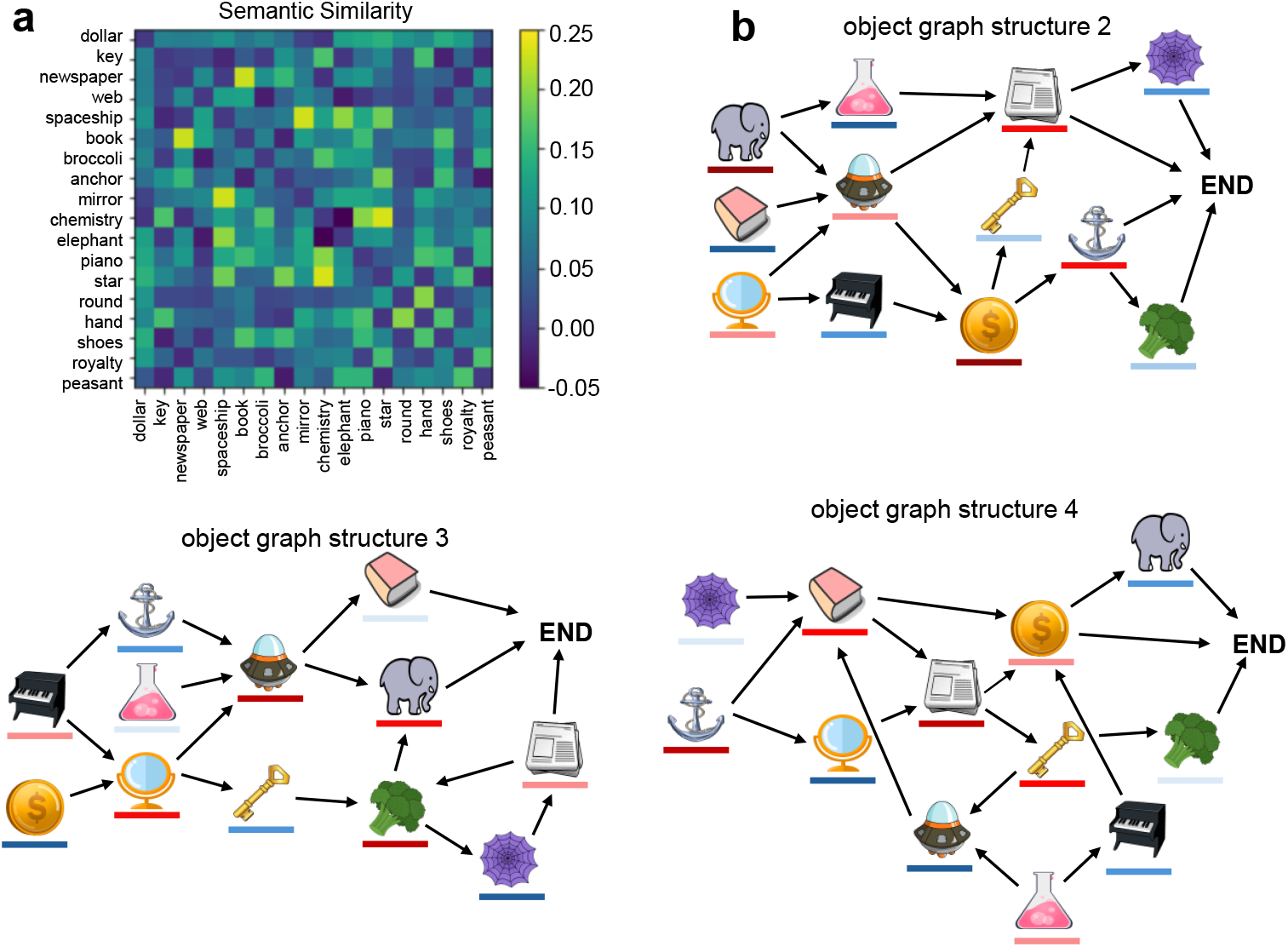
Supplementary task design characteristics. a) Semantic similarity between all used stimuli. Stimuli were chosen with the aim of minimizing semantic correlations between them. B) In addition to the graph structure presented in the main manuscript, three additional graph structures were used.

**Supplementary Figure 2:**
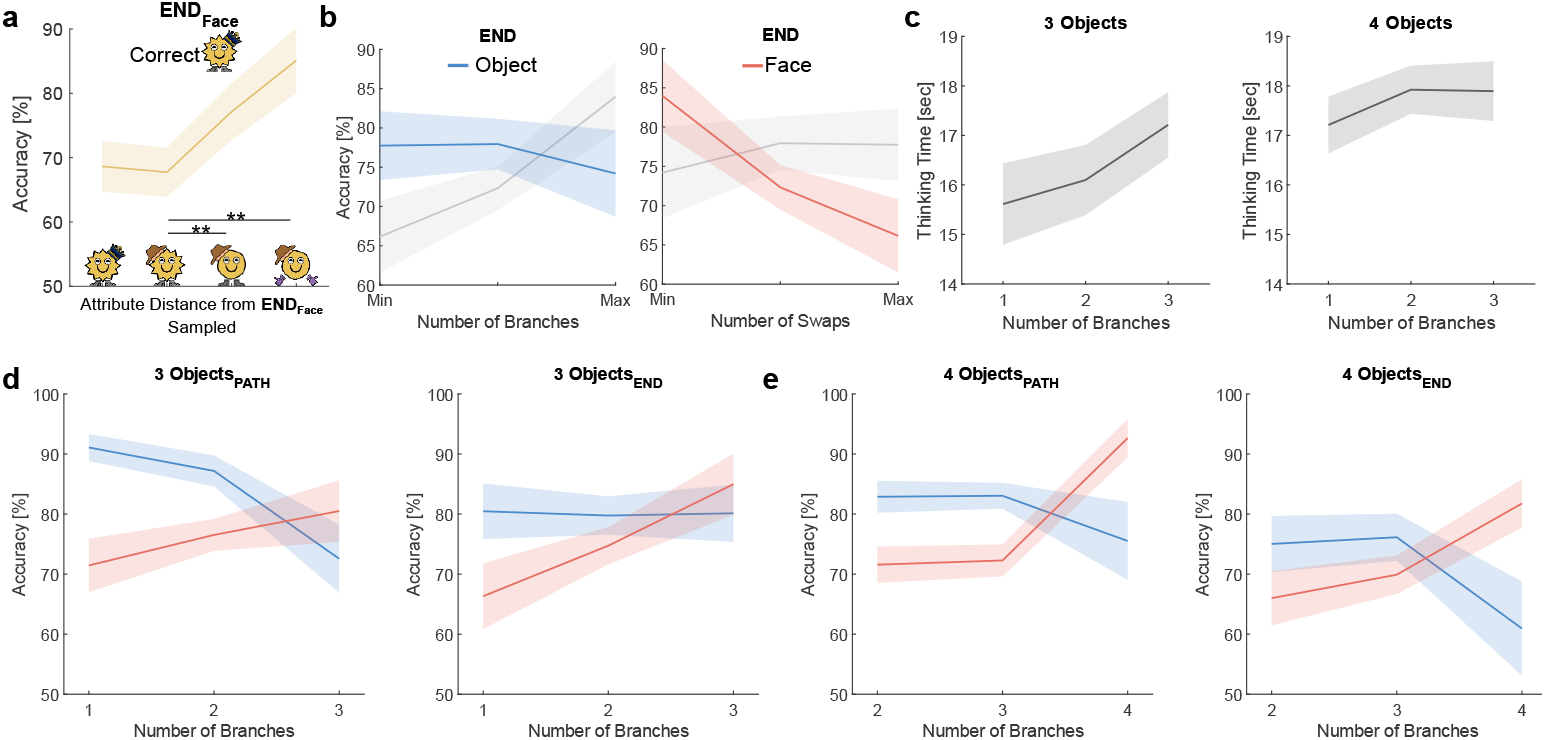
Supplementary behavioral results. a) END Face accuracy increased as a function of the attribute distance between the sampled probe and the target END Face. b) Participants’ accuracy diverged on END Object vs. END Face questions depending on the number of branches on the correct path. c) The pattern in the planning length decrease was consistent when splitting trials based on program length. de) The pattern in diverging accuracy from panel b and Figure 3I was consistent when splitting trials based on program length and question type (PATH, END). Thick lines represent mean accuracy across participants, shaded error bar represents SEM across participants.

**Supplementary Figure 3:**
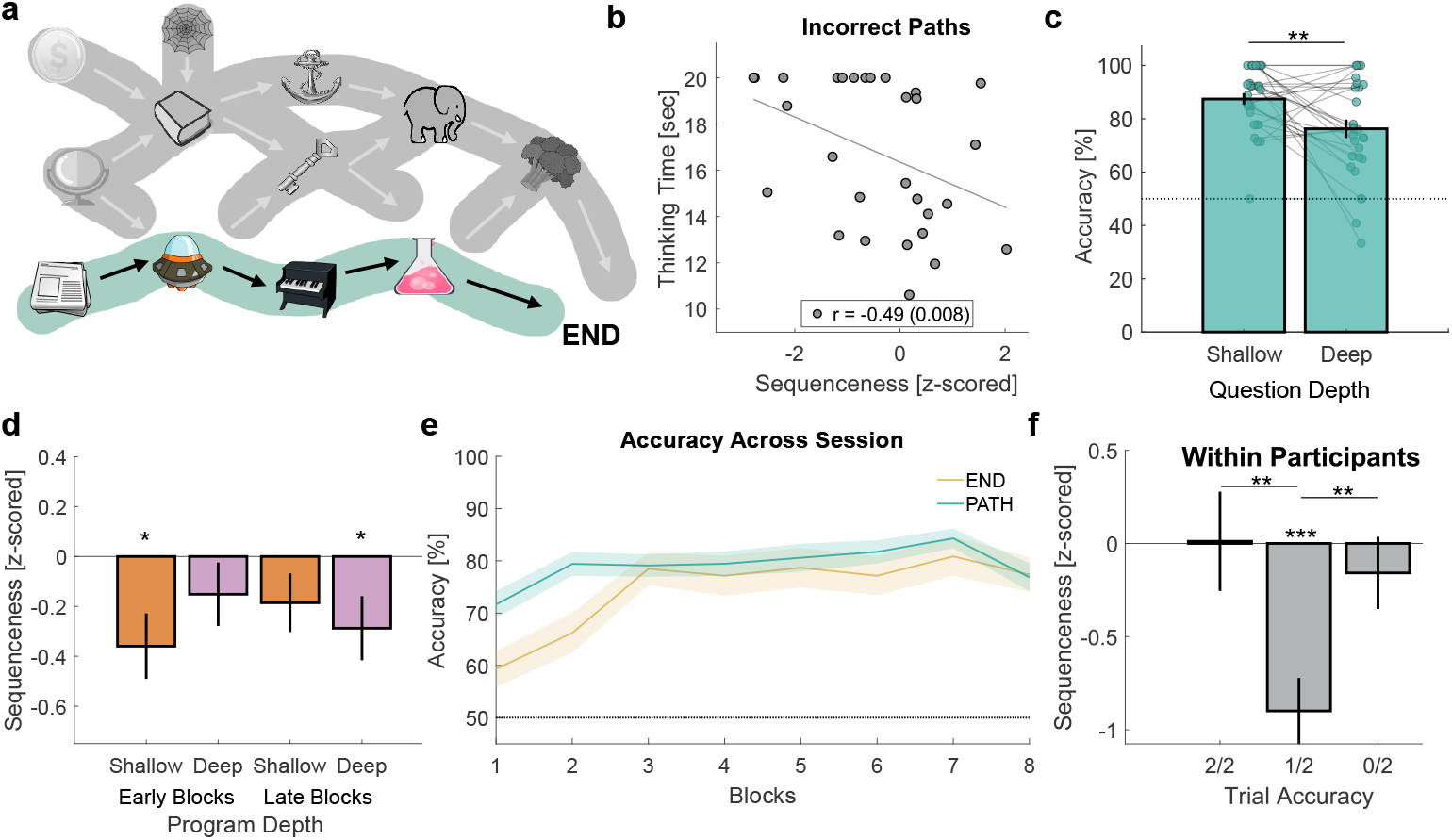
Supplementary object replay-related results. a) Correct paths (green shading) refers to the correct path on a trial based on the starting location and face. Incorrect paths (gray shading) refers to all other paths on the graph on the same trial. b) Correlation between incorrect path replay and participants’ average thinking time. The correlation remained significant after partialling out differences in overall accuracy (r = -.46, p =.014), correct path replay (r = -.49, p =.009), and did not occur due to a correlation between thinking time and ‘replay strength’ (p =.99). c) PATH Accuracy decreased as a function of probe question depth (see Methods: Behavioral analysis). d) Replay peak from Figure 5G split based on the first half (Early) and second half (Late) of the session. e) Participants’ accuracy improved over the duration of the session. f) Incorrect program path replay peak (60 msec, corrected for multiple comparisons) split based on participants’ probe question accuracy. Error bars represent SEM across participants.

**Supplementary Figure 4:**
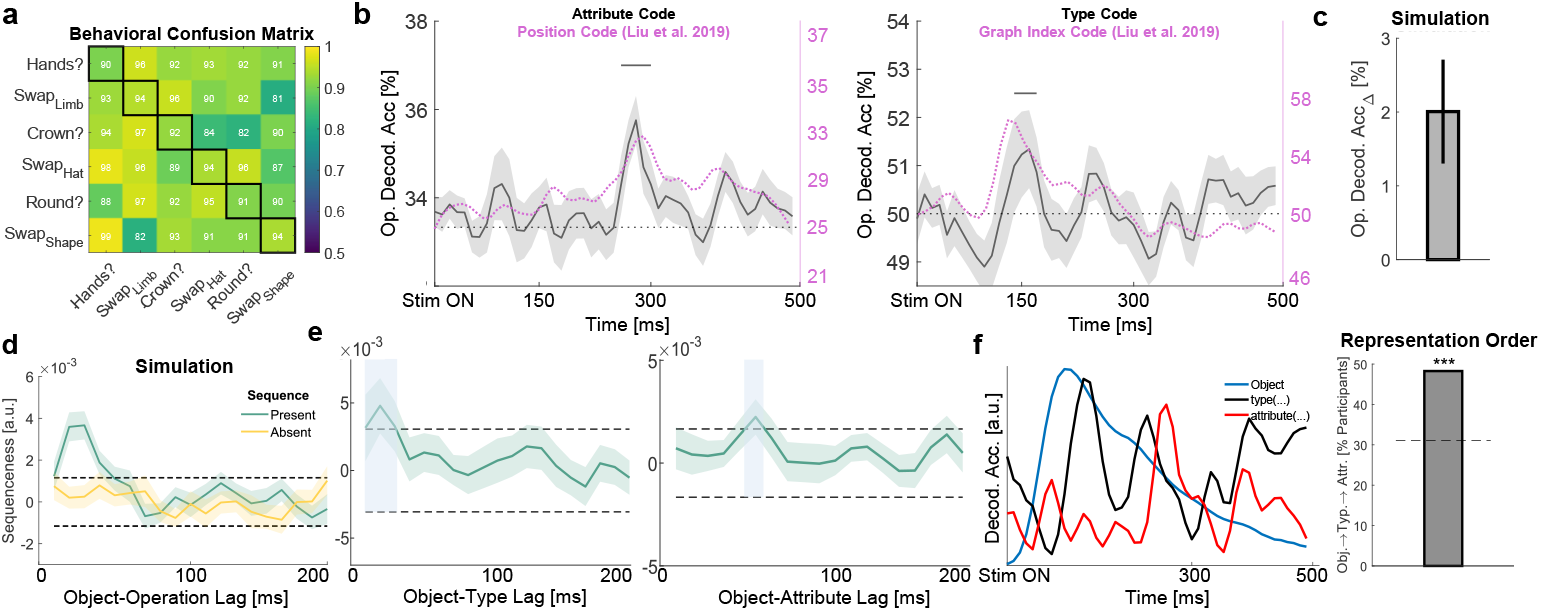
Supplementary operation-related results. a) Behavioral confusion matrix on the operation localizer. Numbers along the diagonal represent the probability of correctly accepting the mapping between an object and subsequent operation text label. Numbers off the diagonal represent the probability of correctly rejecting the mapping between image and subsequent operation text label. b) Operation Attribute (left panel) and Type (right panel) decoding across participants in the operation localizer. Shading represents SEM across participants. The black bar denotes significance in cluster-based permutation testing (exceeding the 97.5th percentile of the mass-corrected null). Overlaid are the decoding results of abstract representations from Liu et al. ^1^. c) Peak operation decoding accuracy of a decoding model generated using synthetic neural data (n = 20 simulated participants). d) The decoding model from panel c was used to generate synthetic operation reactivations. Applying the TDLM analysis distinguished a condition where Location → Operation sequences existed (green line) from a control condition where they did not (yellow line). e) Sequenceness during the reasoning period for Location → Type (left panel) and Location → Attribute (right panel). Dashed lines indicate the 95th percentile of the null distribution (multiple-comparisons corrected). f) The Location → Type → Attribute representational order implied by the reasoning task sequenceness effect was also present within the operation localizer in 13/29 participants (p <.001, Binomial Test, see Methods: Location → Operation sequences). The dashed line represents the 97.5th percentile of a null distribution where representation identity was shuffled.

**Supplementary Figure 5:**
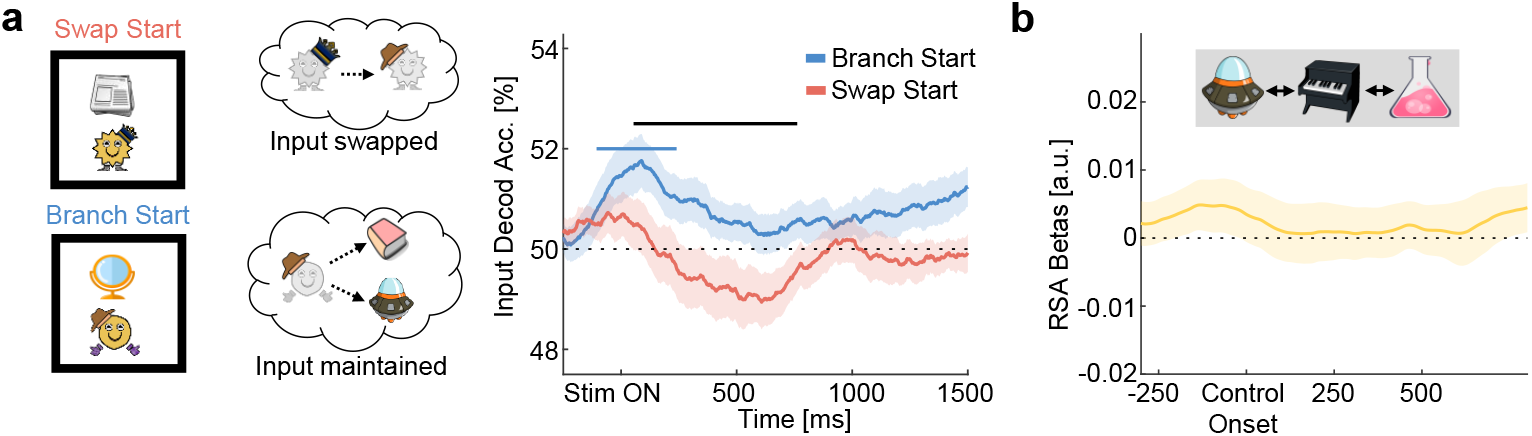
Supplementary program execution-related results. a) Left panel: relationship between input representations and the starting locations’s operation. Swap Starts predict swapping (Crown → Fedora): decreased decoding of the input representation. Branch Starts predict a maintained input representation to move to the next location. Right panel: At Reasoning Start, the input representation was decoded above chance (blue bar) and was stronger on Branch Start compared to Swap Start trials (black bar). Colored bars represent significance in cluster-based permutation testing (exceeding the 97.5th percentile of the mass-corrected null). b) Control analysis for the result in Figure 7H, locked to onsets of control events (see Methods: Replay-aligned RSA analysis). The thick line represents the mean activity across participants, averaged over several control event initializations (n = 5). Shading represents SEM across participants.

